# Structural Landscape of MHC Complexes through Fold Mining

**DOI:** 10.64898/2026.11.23.768646

**Authors:** Giovanna Santos de Lima, Paulo Henrique Alves dos Santos, Samuel Chagas de Assis, Pedro Guilherme Barreto de Santana Nazare, Carlos Daniel Marques Santos Simões, Elton José Ferreira Chaves, Gabriel Ernesto Jara, Rocío Lucía Beatriz Riveros Maidana, Helder Veras Ribeiro-Filho

**Affiliations:** Brazilian Biosciences National Laboratory, Brazilian Center for Research in Energy and Materials, Campinas, São Paulo, Brazil; Graduate Program in Pharmaceutical Sciences, Faculty of Pharmaceutical Sciences, University of Campinas, Campinas, São Paulo, Brazil; Department of Fundamental Chemistry, Center for Natural and Exact Sciences, Federal University of Pernambuco, Recife, Pernambuco, Brazil

**Keywords:** MHC, protein fold, immune recognition

## Abstract

Major histocompatibility complex (MHC) proteins share a characteristic structural platform formed by two α-helices positioned above a β-sheet (α1–α2 domain), which can accommodate ligands and provide an exposed surface for recognition by protein binding partners. This interaction system is best characterized for classical MHC-I molecules presenting peptides to T-cell receptors (TCRs), one of the most extensively studied mechanisms of immune recognition. However, non-classical MHC and MHC-like proteins reveal that the α1–α2 platform is considerably more versatile, supporting a remarkably diverse range of biological functions. Related architectures are also exploited by pathogens, further expanding the diversity of molecular interactions associated with this fold. Yet how this functional diversity relates to structural variation within the shared α1–α2 platform and to the diverse modes of recognition employed by natural and engineered binding partners has not been systematically examined across the broader MHC fold. Here, we systematically mapped MHC α1–α2-like domains across experimentally determined structures in the Protein Data Bank and predicted structures in the AlphaFold Protein Structure Database using Foldseek, and compiled their natural and engineered binding partners. Our comparative analyses revealed substantial heterogeneity across multiple structural layers, including α1–α2 geometry, ligand-binding cavity architecture, and the physicochemical properties and spatial conservation of exposed recognition surfaces, despite the overall conservation of the fold. We further characterized a diverse repertoire of natural and engineered binding partners, revealing distinct structural architectures, interface compositions, and recognition geometries. Together, these findings provide an integrated structural landscape connecting the diversity of the MHC fold with the multiple modes of molecular recognition it supports, offering a comparative framework for understanding these interactions and guiding the engineering of novel MHC-targeting proteins.

## 1 Introduction

The major histocompatibility complex (MHC) encompasses one of the most diverse molecular recognition systems in biology. In humans, the MHC is located on chromosome 6 and comprises a multigene locus conventionally organized into class I, class II, and class III regions. The first two contain genes encoding MHC class I (MHC-I) and MHC class II (MHC-II) molecules, respectively, whereas the class III region includes genes involved in immune and inflammatory responses [1, 2, 3]. Classical MHC-I molecules are expressed by virtually all nucleated cells [4] and, in humans, are represented by the highly polymorphic human leukocyte antigen (HLA)-A, HLA-B, and HLA-C molecules [5]. Together with MHC-II, these proteins mediate antigen surveillance by displaying peptide fragments for recognition by T-cell receptors (TCRs), thereby playing a central role in adaptive immunity [6].

Underlying classical MHC-I antigen presentation is a characteristic molecular architecture. The extracellular region of classical MHC-I (class Ia) consists of a heavy chain containing the α1, α2, and immunoglobulin-like α3 domains, non-covalently associated with β2-microglobulin (β2m). The α1 and α2 domains form a characteristic platform in which two semi-parallel α-helices rest above an antiparallel β-sheet, generating the peptide-binding groove [2, 3]. This groove typically accommodates peptides of 8–10 amino acids derived from intracellular proteins [7]. Peptide loading and editing are supported by accessory proteins, including tapasin (TAPBP), a component of the MHC-I peptide-loading complex [8]. The resulting peptide–MHC complex (pMHC) presents a continuous molecular surface in which exposed peptide residues and the surrounding MHC helices are simultaneously recognized by TCRs [9, 7]. Extensive sequence polymorphism, particularly among residues shaping the peptide-binding groove and its exposed surface, enables different MHC alleles to accommodate distinct peptide repertoires while maintaining the overall α1–α2 architecture [10]. Consequently, substantial chemical and sequence diversity can be displayed within a shared structural framework, generating an exceptionally diverse system of molecular recognition.

Classical peptide presentation to TCRs, however, represents only one biological implementation of this architecture. Closely related non-classical human MHC-I molecules (class Ib), including HLA-E, HLA-F, and HLA-G, preserve the general MHC-I organization but display more restricted polymorphism, distinct expression patterns, and specialized ligand and receptor repertoires [1, 2]. HLA-E presents a relatively restricted peptide repertoire and can be recognized by both TCRs and CD94/NKG2 receptors [11, 12], thereby connecting antigen presentation with the regulation of natural killer (NK) and T-cell responses. HLA-G similarly participates in immune regulation through interactions with members of the leukocyte immunoglobulin-like receptor (LILR) and killer-cell immunoglobulin-like receptor (KIR) families [13, 14, 15]. HLA-F, in turn, can exist as peptide-loaded complexes or peptide-free open conformers, which exhibit distinct recognition patterns involving members of the LILR and KIR receptor families [16, 17, 18]. Thus, even among molecules closely related to classical MHC-I, a common structural framework supports substantially different modes of immune recognition.

The functional breadth associated with this architecture becomes considerably greater when the broader collection of MHC-I-like proteins is considered. These molecules share the characteristic α1–α2 structural platform to varying degrees but can differ substantially in sequence, domain organization, ligand specificity, cellular localization, and biological function [2, 3]. Rather than being restricted to peptides, their internal cavities can accommodate chemically distinct classes of molecules. CD1 proteins, for example, contain predominantly hydrophobic cavities specialized for lipid binding. In humans, the CD1 family comprises CD1a, CD1b, CD1c, CD1d, and CD1e, with CD1a–c classified as group I, CD1d as group II, and CD1e as group III [19]. While CD1a–d present lipid antigens to T cells, CD1e primarily functions intracellularly in lipid processing and transfer [20]. Together, these proteins participate in the recognition of a broad repertoire of self, environmental, and microbial lipids, extending MHC-associated antigen presentation beyond peptides. MR1 provides another striking variation on this molecular architecture. Rather than peptides or lipids, MR1 binds small organic molecules and presents selected metabolites to MR1-restricted T cells, most prominently mucosal-associated invariant T (MAIT) cells. The best-characterized antigens are intermediates derived from microbial riboflavin biosynthesis [21], but the repertoire of MR1-binding molecules also includes other microbial and host-associated metabolites, drug metabolites, and synthetic drug-like compounds [22, 23]. Thus, peptide, lipid, and metabolite recognition can all be supported by variations of the same general MHC-like structural framework.

Importantly, the presence of an MHC-like α1–α2 architecture does not necessarily imply antigen or ligand presentation. Several MHC-like proteins retain, remodel, or substantially reduce the canonical groove while functioning predominantly through interactions with macromolecular binding partners. Endothelial protein C receptor (EPCR), for example, contains a hydrophobic cavity occupied by phospholipids and interacts with the Gla domains of protein C and coagulation factor VII (F7), participating in the regulation of coagulation-related pathways [24, 25]. The hereditary hemochromatosis protein HFE (originally known as HLA-H) retains an MHC-I-like architecture and participates in iron homeostasis through interactions with transferrin receptor 1 (TfR1) [26]. Similarly, the neonatal Fc receptor (FcRn) uses an MHC-I-like scaffold to bind immunoglobulin G (IgG) and albumin in a pH-dependent manner, regulating their intracellular trafficking, transcytosis, and protection from lysosomal degradation [27, 3]. MHC class I chain-related proteins A and B (MICA and MICB), together with members of the UL16-binding protein (ULBP) family, are stress-associated cell-surface proteins that lack conventional antigen-presenting functions and interact with the activating immunoreceptor NKG2D [2, 28]. Zinc-α2-glycoprotein (ZAG) is a soluble MHC-I-like protein associated with lipid metabolism, with reported protein-binding partners including prolactin-inducible protein (PIP) and amine oxidase copper-containing 3 (AOC3) [2, 29, 30]. However, no physiological protein-binding partner has yet been structurally characterized engaging the upper α1–α2 recognition surface.

The versatility of this architecture is not restricted to human endogenous proteins. Pathogens, particularly viruses, also exploit the MHC fold through diverse strategies. One strategy involves molecular mimicry of MHC-like α1–α2 domains to engage receptors that normally recognize host MHC-like proteins. For example, the secreted orthopoxvirus protein OMCP adopts an MHC-I-like platform and binds NKG2D, competitively blocking recognition of its natural binding partners [31], whereas the human cytomegalovirus (HCMV) MHC-I homolog UL18 binds leukocyte immunoglobulin-like receptor B1 (LILRB1) [32, 33]. Viral proteins can also repurpose the MHC fold toward different recognition systems, as exemplified by the poxvirus protein 2L, which adopts an MHC-I-like architecture but binds tumor necrosis factor α (TNF-α) and prevents its interaction with TNF receptors [34]. Another strategy involves directly targeting host MHC and MHC-like molecules to alter their trafficking or promote degradation. Examples include adenovirus E3-19K, which retains classical MHC-I molecules in the endoplasmic reticulum [35, 36]; HCMV US2, which targets MHC-I for proteasomal degradation [37]; and HCMV UL16, which promotes the intracellular sequestration of MICB and selected ULBP proteins [38]. Conversely, viruses can exploit MHC-like proteins as cellular receptors, as demonstrated for the interaction of FcRn with echovirus capsid proteins [39, 40] and human astrovirus spike proteins (VP27, derived from the capsid precursor VP90) [41, 42]. Similar strategies extend beyond viruses; for example, Plasmodium falciparum uses the cysteine-rich interdomain region α1 (CIDRα1) of P. falciparum erythrocyte membrane protein 1 (PfEMP1) to bind EPCR on endothelial cells, promoting cytoadhesion while interfering with the physiological interaction of EPCR with protein C [43, 44].

Given the biological relevance of MHC recognition interfaces, considerable efforts have been directed toward their modulation using engineered binding proteins. Antibodies and antibody-derived formats, including single-domain antibodies (VHHs) and single-chain variable fragments (scFvs), have been developed to mimic TCR recognition of peptide–MHC complexes [45, 46] or to block native protein interactions, as exemplified by anti-FcRn antibodies that prevent IgG binding and recycling [47]. More recently, advances in computational protein design and artificial intelligence have enabled the development of non-TCR scaffolds capable of highly specific recognition of peptide–MHC-I complexes, including *de novo* designed miniproteins and α-helical binding modules [48, 49]. Alternative engineering strategies have also produced peptide-dependent MHC binders based on repurposed protein scaffolds, such as TRACeR-I [50], and designed ankyrin repeat proteins (DARPins) targeting specific peptide–MHC complexes [51]. Together, these approaches illustrate the growing diversity of engineered protein architectures capable of recognizing and modulating MHC-associated interactions.

Together, these examples illustrate that the MHC fold represents a remarkably versatile molecular recognition scaffold whose functional repertoire extends well beyond canonical peptide presentation and immune recognition. Despite decades of structural studies on individual MHC and MHC-like families and the increasing availability of experimentally determined complexes, an integrated comparative characterization linking the structural and physicochemical diversity of the MHC fold to the architecture and recognition modes of its binding partners remains limited. Such a framework could help identify common and family-specific features of molecular recognition, reveal the range of structural solutions used by different binding partners, and provide insights for the engineering of proteins capable of modulating MHC-associated interactions.

Here, we present a systematic structural analysis of the MHC fold across its broad functional diversity. By mining the Protein Data Bank (PDB) and AlphaFold Protein Structure Database (AFDB) using Foldseek, we assembled a dataset of MHC-fold proteins and their experimentally characterized complexes with natural and engineered binding partners. We then compared multiple complementary dimensions of this molecular recognition platform, including global structural similarity, α1–α2 geometry, ligand-binding cavity architecture and occupation, physicochemical properties of exposed surfaces, and the structural and chemical characteristics of MHC–binder interfaces. We also characterized the architectures and recognition modes of the binding partners themselves. By integrating these properties across classical and non-classical MHC molecules, MHC-like families, pathogen-derived proteins, and engineered recognition systems, we provide a comparative structural framework for understanding the versatility of the MHC fold as a molecular recognition platform.

## 2 Results

### 2.1 Structural diversity of MHC-fold proteins and their binding partners

By mining the PDB using Foldseek, we identified a total of 1,267 entries containing domains homologous to the MHC-I α1–α2 platform, considering only human-derived proteins or viral proteins that biologically interact with human proteins. Mining of the AlphaFold Database (AFDB) identified 526 entries after filtering based on sequence length and taxonomic origin, 25 of which corresponded to reviewed UniProt entries. The proteins identified across both datasets encompassed the four major groups of MHC-fold proteins: classical MHC molecules (HLA-A, HLA-B, and HLA-C), non-classical MHC molecules (HLA-E, HLA-F, and HLA-G), MHC-like proteins (including the CD1, MR1, FcRn, EPCR, MIC, ZAG, HFE, and ULBP families), and viral MHC mimetics (including UL18, OMCP, and 2L) (Figure 1).

**Figure 1:**
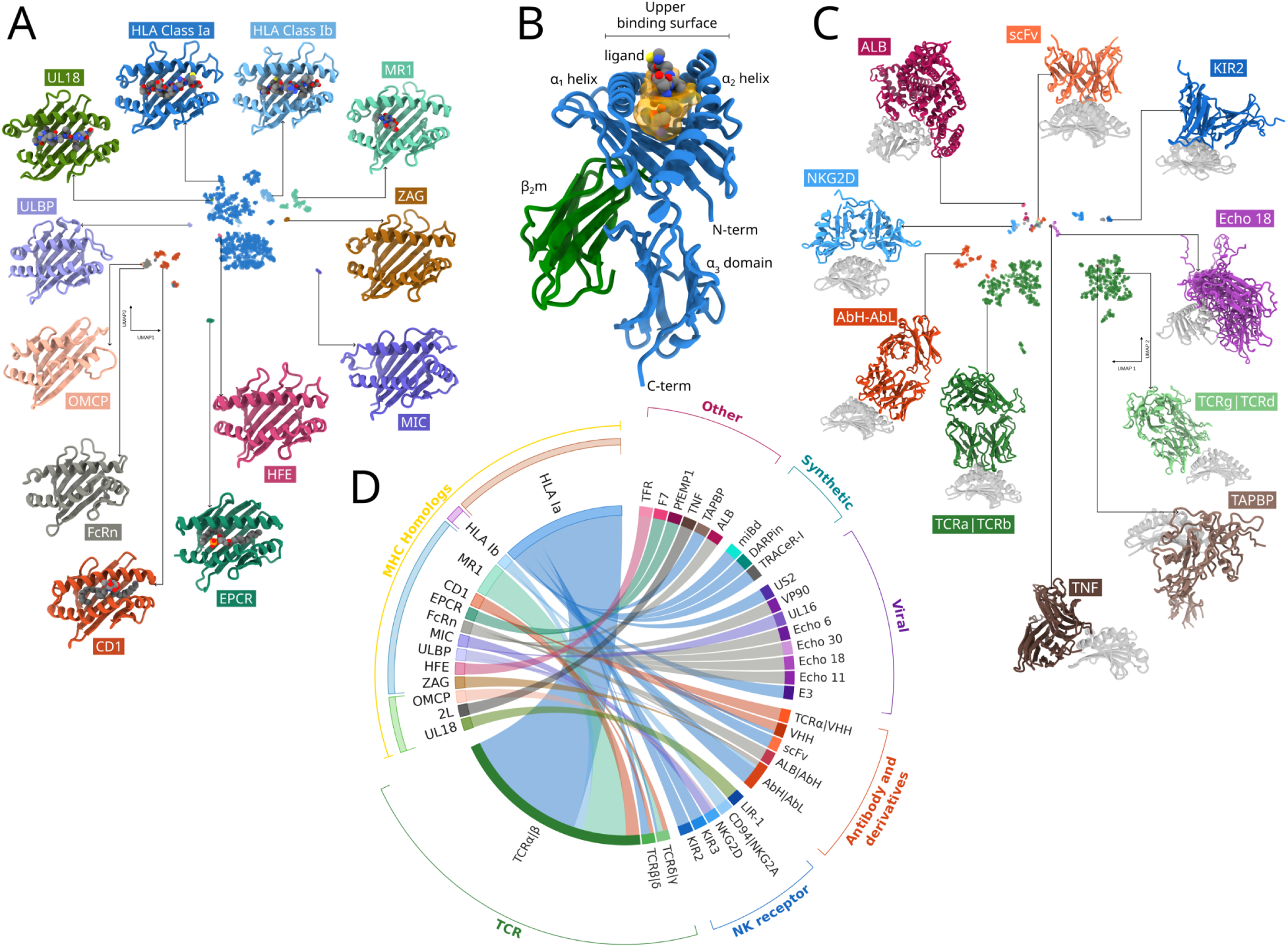
Structural landscape of MHC-fold proteins and their binding partners derived from structural database mining. (A) Two-dimensional UMAP representation of MHC-fold proteins identified in the PDB, generated from pairwise TM-score similarities of their α1–α2 domains. The UMAP was calculated using the combined PDB and AFDB dataset, but AlphaFold models, single-chain (SC), and chimeric constructs were omitted from this panel for visual clarity, while preserving the original UMAP coordinates (see Supplementary Figure S1A for the complete representation). Structures are colored according to MHC family, with representative structures shown for selected groups. (B) Representative MHC-I structure (PDB ID: 1A1M) highlighting the α1 and α2 helices, the ligand-binding groove, β2-microglobulin (β2m), the α3 domain, and the upper α1–α2 recognition surface considered for the analysis of MHC-binding partners. (C) UMAP representation of the structural diversity of MHC-binding partners identified in the PDB, based on pairwise TM-score similarities. Representative binders from selected structural groups are shown in color, with their interacting MHC proteins shown in gray. (D) Chord diagram summarizing structurally characterized interactions between MHC-fold protein families and their macromolecular binding partners identified in the PDB. MHC proteins are grouped on the left, while binders are organized according to major structural or functional classes on the right. Connecting ribbons represent MHC–binder associations observed in experimentally determined complexes.

Thus, all currently recognized families of human MHC-fold proteins are represented among the available experimental structures, with at least one structure from each family and multiple allelic variants of polymorphic HLA proteins. The PDB dataset was particularly enriched in classical MHC molecules, which accounted for 79% of the identified entries. Within the classical MHC-I dataset, HLA-A was more extensively represented than HLA-B and HLA-C, with a particularly strong enrichment for the HLA-A*02 allele group, which accounted for 68% of HLA-A entries (Supplementary Figure S2). To account for potential overrepresentation of equivalent experimental complexes, redundancy among the processed PDB structures was also evaluated as described in the Methods section. Analyses restricted to nonredundant structures are explicitly indicated in the corresponding results sections. Unresolved internal regions within the MHC *α*_1_–*α*_2_ platform were modeled as also described in the Methods.

Notably, under our structural search and filtering criteria, no previously unrecognized human protein families adopting the MHC fold were identified among the AFDB entries, even when both reviewed and unreviewed UniProt entries were considered. Thus, despite substantially expanding the search space beyond experimentally determined structures, mining the AFDB did not reveal additional human MHC-fold protein families beyond those already represented in the PDB.

Within the PDB dataset, 32% of entries corresponded to fully bound complexes, in which the MHC α1–α2 domains were associated with a ligand occupying the binding groove and a macromolecular binder interacting with the upper helical surface. Most of these entries corresponded to classical MHCs presenting peptides to TCRs, although complexes involving the MHC-like molecules MR1, EPCR, and CD1, as well as the non-classical HLA-E, were also represented. Overall, 94% of entries contained a ligand bound to the MHC groove, whereas most ligand-free structures corresponded to MHC-fold proteins for which no natural groove-binding ligand is known, such as FcRn, MIC, and ULBP. A small fraction (2% of all entries) corresponded to MHC-fold proteins that naturally present ligands, such as classical MHCs, CD1, MR1, EPCR, and ZAG, but whose structures were determined in a ligand-unbound state. Finally, 36% of all entries contained a macromolecular binder interacting with the MHC α1–α2 domains.

In addition to accommodating ligands, MHC-fold proteins can also undergo post-translational N-linked glycosylation. Within the PDB dataset, 85 structures contained at least one experimentally resolved N-linked glycosylation site, totaling 159 glycosylation instances. Mapping these sites onto the corresponding gene-specific reference structures revealed 30 distinct glycosylation sites across 13 proteins, of which 6 were located within the *α*_1_ helix, 2 within the *α*_2_ helix, and 22 outside the helical regions (Supplementary Figure S3). Although most glycosylation sites were located outside the two helices, their spatial distribution varied across MHC-fold proteins, even among members of the same protein family. Glycosylation sites along the *α*_1_ helix were observed in CD1a, CD1b, CD1c, 2L, and ULBP, whereas sites along the *α*_2_ helix were identified in CD1d and EPCR. Importantly, these findings are restricted to experimentally resolved glycans in the available PDB structures and may not fully reflect the diversity of N-linked glycosylation sites across MHC-fold proteins.

A Uniform Manifold Approximation and Projection (UMAP) representation based on pairwise structural similarity (TM-score) among all identified MHC-fold protein structures revealed an organized structural landscape, with clusters largely corresponding to individual MHC families (Figure 1 and Supplementary Figure S1). Classical MHC-I molecules formed the largest cluster, predominantly comprising HLA-A and HLA-B alleles, reflecting both their structural conservation and their extensive representation in the PDB. Non-classical MHC molecules occupied regions adjacent to classical MHC-I, with HLA-F and HLA-G particularly close to this group (Supplementary Figure S1). Pairwise comparisons between representative structures supported these relationships, with high TM-scores and low RMSD values demonstrating strong conservation of the α1–α2 architecture and providing a direct quantitative measure of structural similarity between families (Supplementary Figure S4). The few subclusters observed within classical MHCs, particularly among HLA-B structures, were primarily associated with local conformational variations in coil regions connecting the β-sheet strands rather than major rearrangements of the overall fold.

MHC-like proteins also displayed a pronounced family-dependent structural organization, with most families forming well-defined clusters in the UMAP representation (Figure 1A). This organization was supported by pairwise structural comparisons of representative structures, which revealed high within-family similarity together with variable degrees of divergence between families (Supplementary Figure S4). Among the MHC-like proteins, MR1, ZAG, HFE, and the viral MHC mimic UL18 were among the most structurally similar to classical and non-classical MHC-I, generally displaying representative TM-scores above 0.8 and defining an MHC-I-related region of the structural landscape. Particularly, MR1 retained very high similarity to classical MHC-I despite its distinct ligand specificity, with representative TM-scores of approximately 0.90. CD1 molecules occupied a separate region and formed isoform-specific subclusters in the UMAP representation. Nevertheless, comparisons among CD1 isoforms yielded TM-scores above 0.9, indicating strong conservation of a characteristic CD1 α1–α2 architecture despite subtle isoform-specific differences. EPCR and FcRn occupied additional defined regions of the structural landscape, while MIC and ULBP formed distinct clusters, further illustrating the emergence of family-specific variations of the MHC fold.

Viral MHC mimics were particularly informative regarding the structural plasticity of the fold. In contrast to UL18, which retains an α1–α2 architecture closely related to classical MHC-I, OMCP and 2L were among the most structurally divergent homologs, exhibiting generally lower TM-scores and higher RMSD values relative to the other MHC families. Nevertheless, their closest structural relationships among the representative structures were observed between OMCP and FcRn (TM-score = 0.73) and between 2L and CD1b (TM-score = 0.76). These relationships are consistent with their relative positions in the UMAP representation, while emphasizing that even highly divergent viral mimics retain recognizable elements of the MHC structural framework.

AlphaFold models closely recapitulated the experimentally determined structures within their respective families, despite being predicted in the absence of their biological ligands. Across all families, structural deviations relative to the best-matching experimental structures remained small, with the largest RMSD values ( 2 Å) observed for ULBP proteins and the lowest deviations observed for classical MHC molecules (Supplementary Figure S5). All MHC-fold protein families also showed high AlphaFold confidence, with mean pLDDT values consistently above 90, including ULBP, which displayed the lowest confidence among the analyzed families (Supplementary Figure S6). These results indicate that AlphaFold reproduces the overall architecture of the MHC fold with high confidence across the analyzed families. This assessment, however, is restricted to the MHC α1 and α2 domains themselves and does not extend to bound ligands or receptor interactions. Furthermore, these confidence values should be interpreted with caution, as AlphaFold predictions may benefit from the presence of homologous experimental structures in its training data.

Experimentally determined single-chain fusion constructs, in which the ligand, MHC molecule, or β2m are covalently linked, were identified for HLA-A and HLA-B (17 structures), MR1 (1 structure), and CD1c and CD1e (5 structures). Chimeric constructs, in which domains from different species are combined, were also identified in the analyzed dataset (4 structures for CD1c and 2 for CD1d). Despite their artificial architecture, these constructs clustered within their respective native families and showed low structural deviations relative to their closest native counterparts, typically below 1 Å RMSD (Supplementary Figure S5). These observations indicate that covalent fusion does not substantially perturb the overall α1–α2 domain fold. Nevertheless, to avoid potential biases associated with their engineered architecture, single-chain or chimeric constructs were excluded from subsequent analyses. Comparisons between experimentally determined ligand-free MHC structures and their best-matching ligand-bound counterparts also revealed high structural similarity of the *α*_1_–*α*_2_ platform, with TM-scores approaching 1 and low RMSD values (Supplementary Figure S5C). These observations indicate that the overall architecture of the MHC fold is largely conserved across the ligand-binding states represented in the available experimental structures.

From the binder perspective, the α1–α2 domains of nearly all analyzed MHC-fold protein families were represented in complexes with at least one macromolecular binding partner. The exceptions were ZAG, for which no physiological receptor engaging the α1–α2 domain is currently known and only an antibody-derived complex was identified in the PDB, and UL18. Although UL18 interacts with leukocyte immunoglobulin-like receptors (LILRs), these receptors do not engage the canonical upper α1–α2 surface considered in our analysis and were therefore excluded from the binder dataset. Despite sharing a common structural scaffold, classical, non-classical, and MHC-like molecules associate with a remarkably broad repertoire of binding partners (Figure 1D), highlighting the versatility of the MHC fold as a molecular recognition platform.

Currently, T cell receptors (TCRs) constitute the most abundant class of MHC receptors in the PDB, reflecting the extensive structural characterization of TCR-mediated recognition of peptide–MHC complexes (Figure 1). A UMAP representation based on pairwise TM-scores revealed well-defined structural clusters corresponding to the major binder families (Figure 1C). TCRα and TCRδ clustered together, as did TCRβ and TCRγ, reflecting their close structural relationships. Antibody light (AbL) and heavy (AbH) chains also formed distinct clusters. Although all of these proteins belong to the immunoglobulin (Ig) superfamily, they differ in the relative orientation and organization of their variable and constant domains. Similarly, TAPBP, despite possessing an overall fold distinct from TCRs and antibodies, clustered in close proximity owing to its conserved Ig-like domain.

An additional cluster was enriched in Ig-like domains represented solely by their variable or MHC-interacting regions, including TCRα, TCRβ, AbH, AbL, VHH, scFv, UL16, and US2. Members of the KIR family also formed distinct clusters, with KIR2 and KIR3 separating according to their two- and three-Ig-domain architectures, respectively. In contrast, receptors lacking Ig-like domains occupied independent regions of the structural landscape. These included the C-type lectin receptors CD94 and NKG2, characterized by mixed α-helical and β-strand architectures; predominantly α-helical binders such as the CIDRα domain of PfEMP1 and albumin; and other structurally distinct binders, including the transferrin receptor (TfR) and Factor VII light chain (F7). α-helical designed miniproteins, including AI-based designed miniprotein binders (miBd) and TRACeR-I, clustered in proximity to the naturally occurring α-helical binders. Finally, a distinct cluster characterized by β-sandwich folds with connecting loops included TNF-family members and echovirus capsid proteins.

Collectively, these analyses reveal a broad structural landscape of proteins sharing the MHC α1–α2 architecture, as represented by experimentally determined structures in the PDB and predicted models in the AFDB. Within the experimentally characterized PDB complexes, these proteins are recognized by a diverse repertoire of structurally distinct binding partners. Importantly, this diversity is not limited to differences between MHC families, as structurally unrelated binders can recognize members of the same MHC family (1D). Classical MHC-I molecules, for example, have been structurally characterized in complexes with TCRs, antibodies, KIRs, viral immunoevasins such as E3-19K and US2, and engineered binders. Similarly, CD1 molecules are recognized by both TCRs and VHH antibodies, whereas MIC proteins interact with NKG2D and the viral protein UL16. EPCR provides another example, engaging structurally distinct partners such as PfEMP1 CIDRα domains and Factor VII. Together, these examples highlight the versatility of the MHC fold as a molecular recognition platform, in which related structural architectures accommodate diverse binding partners and modes of recognition.

### 2.2 Architecture of the α1–α2 platform and ligand-binding cavities across MHC-fold proteins

All analyzed MHC-fold proteins share a common architectural feature: two general α-helice segments positioned along opposite sides of a β-sheet platform, forming an exposed surface for macromolecular recognition and, when present, delimiting a central ligand-binding groove (Figure 1B). Across the analyzed MHC families, the α-helical content of the α1–α2 region varied only modestly, averaging approximately 30–40% (Supplementary Figure S7), underscoring the conservation of this structural element across the MHC fold. Among all analyzed structures, the viral MHC mimetics 2L and OMCP exhibited the lowest α-helical content.

Analysis of secondary-structure profiles further revealed differences in the organization and continuity of the two canonical groove helices across MHC-fold proteins (Supplementary Figure S8). In classical MHC molecules, these helices exhibited a conserved organization comprising four helical subregions (α1N, α1C, α2N, and α2C), a pattern broadly maintained across several MHC-like families. However, notable variations were observed, including a marked reduction of the short N-terminal α1 helical segment in EPCR. Viral MHC mimetics, particularly OMCP and 2L, displayed more divergent profiles, characterized by shortened helical segments and extended non-helical interruptions. In OMCP, both α1 and α2 exhibited substantial discontinuities, whereas 2L showed a pronounced reduction of the α2N helical segment while retaining a longer α2C region.

The relative openness of the α1–α2 platform, measured as the average minimum helix-to-helix distance (see Methods), was 12.7 Å for classical MHC-I and 12.6 Å for non-classical MHC-I molecules (Figure 2), with a relatively narrow distribution despite the extensive allelic diversity represented in the PDB. One upper outlier (9NKH) in the classical MHC-I group appears to result from a structure containing a long unresolved gap. MR1 and the viral MHC mimic UL18, the latter also retaining the ability to present peptides, exhibited similarly large helix separations, consistent with their overall structural similarity to classical MHC-I molecules. This observation is particularly notable for MR1, which retains a TCR-recognized MHC-like platform despite presenting small-molecule metabolites rather than peptides, suggesting that an open α1–α2 geometry can be conserved independently of ligand class. Intermediate distances of approximately 10–12 Å were observed for ZAG, CD1, EPCR, HFE, and MIC, whereas the most compact α1–α2 arrangements were found in FcRn, ULBP, OMCP, and 2L, with mean distances of approximately 8.5–10 Å. These latter molecules lack a canonical antigen-presenting groove, suggesting that compaction of the α1–α2 platform accompanies the repurposing of the MHC fold from ligand presentation toward direct protein–protein recognition. Representative structures illustrating the diversity of cavity geometries and bound ligands across MHC-fold proteins are shown in Figure 2E.

**Figure 2:**
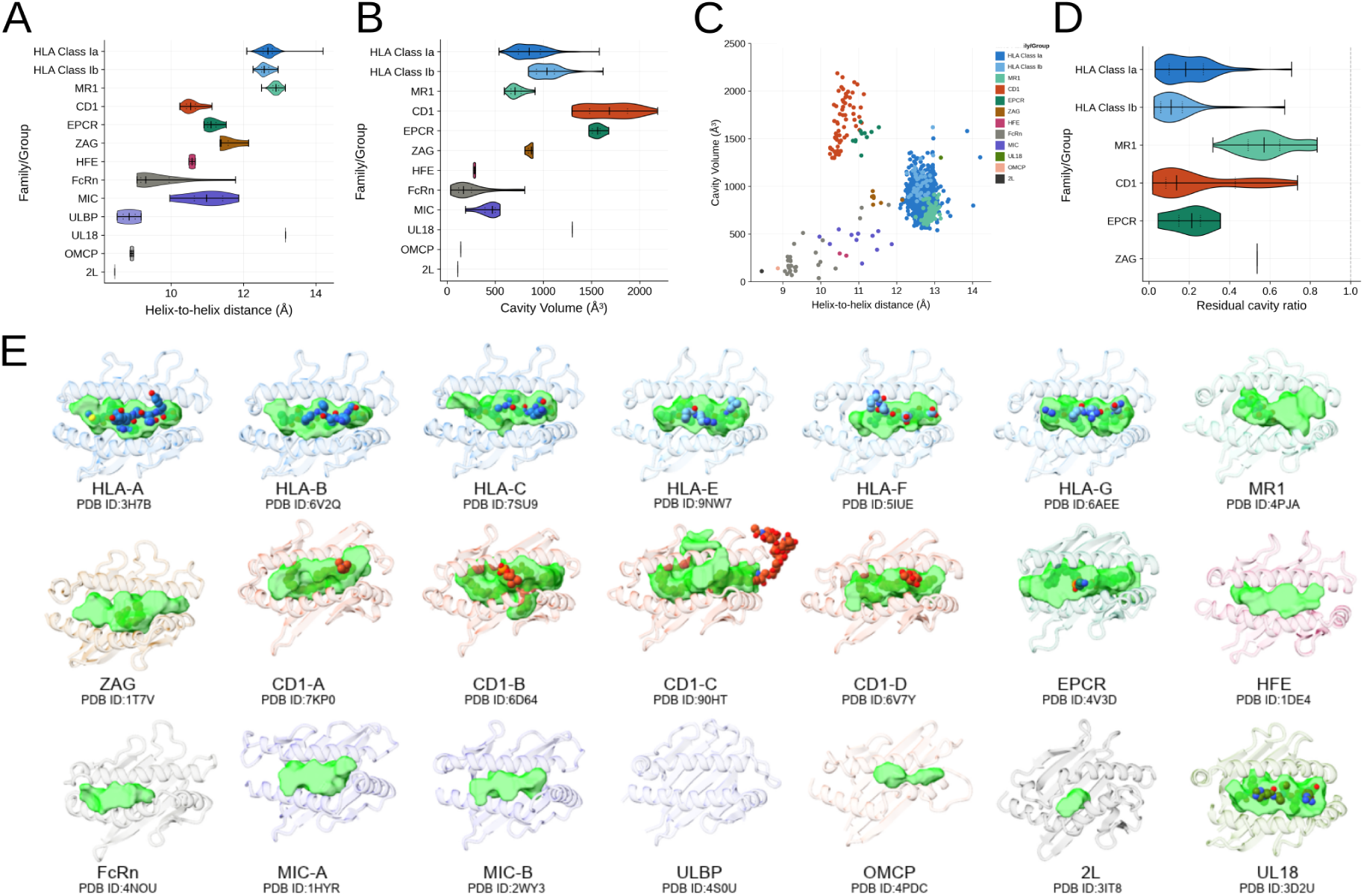
Architecture of the α1–α2 platform and ligand-binding cavities across MHC-fold proteins. (A) Distribution of helix-to-helix distances across MHC-fold protein families, calculated as the mean nearest-neighbor Cα distance between residues of the α1 and α2 helices (see Methods for details). (B) Distribution of cavity volumes across MHC-fold protein families, estimated using pyKVFinder after computational removal of bound ligands, when present. (C) Relationship between helix-to-helix distance and cavity volume for individual structures, colored according to MHC family. (D) Distribution of the residual cavity ratio for ligand-bound structures, calculated as the ratio between the cavity volume detected with the ligand present and that detected after ligand removal. Structures without a detectable residual cavity were excluded from this analysis. (E) Representative structures illustrating the diversity of the α1–α2 platform across MHC-fold proteins, highlighting differences in cavity geometry and helix-to-helix distance. Protein structures are shown as cartoons, detected cavities as green surfaces, and bound ligands as spheres when present. The PDB ID of each representative structure is indicated below the corresponding model. All quantitative analyses were performed using nonredundant structures (see Methods).

The relationship between helix-to-helix distance and cavity volume varied across MHC-fold protein families, rather than following a uniform trend (Figure 2C). Analysis of ligand-binding cavities across the analyzed MHC structures, including naturally unoccupied structures and ligand-bound structures analyzed after computational ligand removal, showed that classical MHC-I molecules have cavity volumes centered at approximately 870 Å³, with most structures ranging from approximately 540 to 1,300 Å³ after excluding outliers associated with the computational modeling of long unresolved regions. HLA-A showed a slightly lower average cavity volume than HLA-B and HLA-C (Supplementary Figure S7E). Non-classical MHC-I molecules displayed slightly larger volumes on average but remained largely within the same range. MR1, which accommodates small-molecule metabolites, showed a smaller average cavity volume ( 720 Å³), although its distribution substantially overlapped that of classical MHC-I. The MHC-like molecule ZAG similarly displayed cavity volumes within the range observed for MHC-I, averaging approximately 860 Å³. Together with their relatively similar α1–α2 helix separations, these results indicate that these molecules retain groove architectures broadly comparable in scale to that of classical MHC-I despite their distinct biological functions and ligand-binding properties.

Groove volume, however, was not determined solely by the separation between the α1 and α2 helices. CD1 and EPCR, both lipid-binding molecules, represented striking examples of this decoupling, displaying substantially shorter helix-to-helix distances than classical MHC-I while forming considerably larger cavities, frequently exceeding 1,500 Å³ (Figure 2 and Supplementary Figure S7). This difference was accompanied by greater cavity depth, indicating that ligand-accommodating space can expand toward the protein interior without a corresponding increase in helix separation, as illustrated by the relationship between helix openness and cavity volume (Figure 2C). Comparison of cavity volumes before and after computational removal of bound ligands further revealed distinct modes of cavity occupation (Figure 2D). Classical and non-classical MHC-I molecules retained only a small fraction of their available cavity in the presence of peptide, indicating extensive occupation of the groove (exceptions reflect modified peptides in classical MHC-I or noncanonical peptide-binding modes, as observed for HLA-F). In contrast, MR1 retained a considerably larger fraction of its cavity when bound to small-molecule metabolites, despite substantial overlap with classical MHC-I in overall cavity volume. CD1 showed a broad distribution of residual cavity fractions, consistent with variable occupation of its exceptionally large and deep cavities by structurally diverse lipid ligands. Together, these results show that helix-to-helix separation, cavity volume and depth, and the fraction of cavity occupied by bound ligands capture complementary aspects of α1–α2 architecture and reveal distinct strategies by which the MHC fold accommodates chemically and structurally diverse ligands.

Several MHC-fold proteins lacking a conventional ligand-presenting function exhibited substantially reduced cavities. MIC, HFE, and FcRn displayed average cavity volumes of 435, 285, and 260 Å³, respectively, while the viral MHC mimics OMCP and 2L exhibited even smaller cavities. No cavity was detected in the ULBP family under the applied detection criteria. More generally, the reduced cavities observed in MIC, HFE, FcRn, OMCP, and 2L, together with the absence of a detectable cavity in ULBP, are consistent with extensive remodeling or loss of the canonical ligand-accommodating architecture in MHC-like scaffolds specialized predominantly for direct protein–protein recognition.

### 2.3 Physicochemical diversity of the MHC α1–α2 recognition surface

Despite the overall structural conservation of the MHC fold, the physicochemical properties of the exposed α1–α2 surface vary across MHC-fold proteins and may contribute to the recognition of distinct molecular partners. To characterize this chemical diversity within the experimentally determined MHC structures available in the PDB, we analyzed the amino-acid composition of solvent-exposed residues within the α1 and α2 helices across MHC families (Figure 3A and Supplementary Figure S9A). Charged residues constituted a substantial fraction of the exposed helical surface in most families, with Glu, Asp, Arg, and Lys frequently among the most abundant residues. Polar uncharged residues, particularly Gln, also made substantial contributions. However, when residues were grouped according to physicochemical classes, the relative contributions of hydrophobic, aromatic, polar uncharged, acidic, and basic residues varied considerably among families. Thus, rather than displaying a uniform physicochemical composition, MHC-fold proteins exhibited distinct balances among these residue classes. Cys was nearly absent from the analyzed surfaces, whereas Phe, Trp, Pro, and His were generally less frequent, with family-specific exceptions. Electrostatic potential maps calculated for selected MHC structures further revealed how differences in amino-acid composition translate into distinct spatial distributions of electrostatic potential across the α1–α2 surface (Figure 3C). Although most MHC proteins exhibited heterogeneous distributions of positive, negative, and near-neutral regions, their relative extent and spatial organization varied considerably among the analyzed proteins.

**Figure 3:**
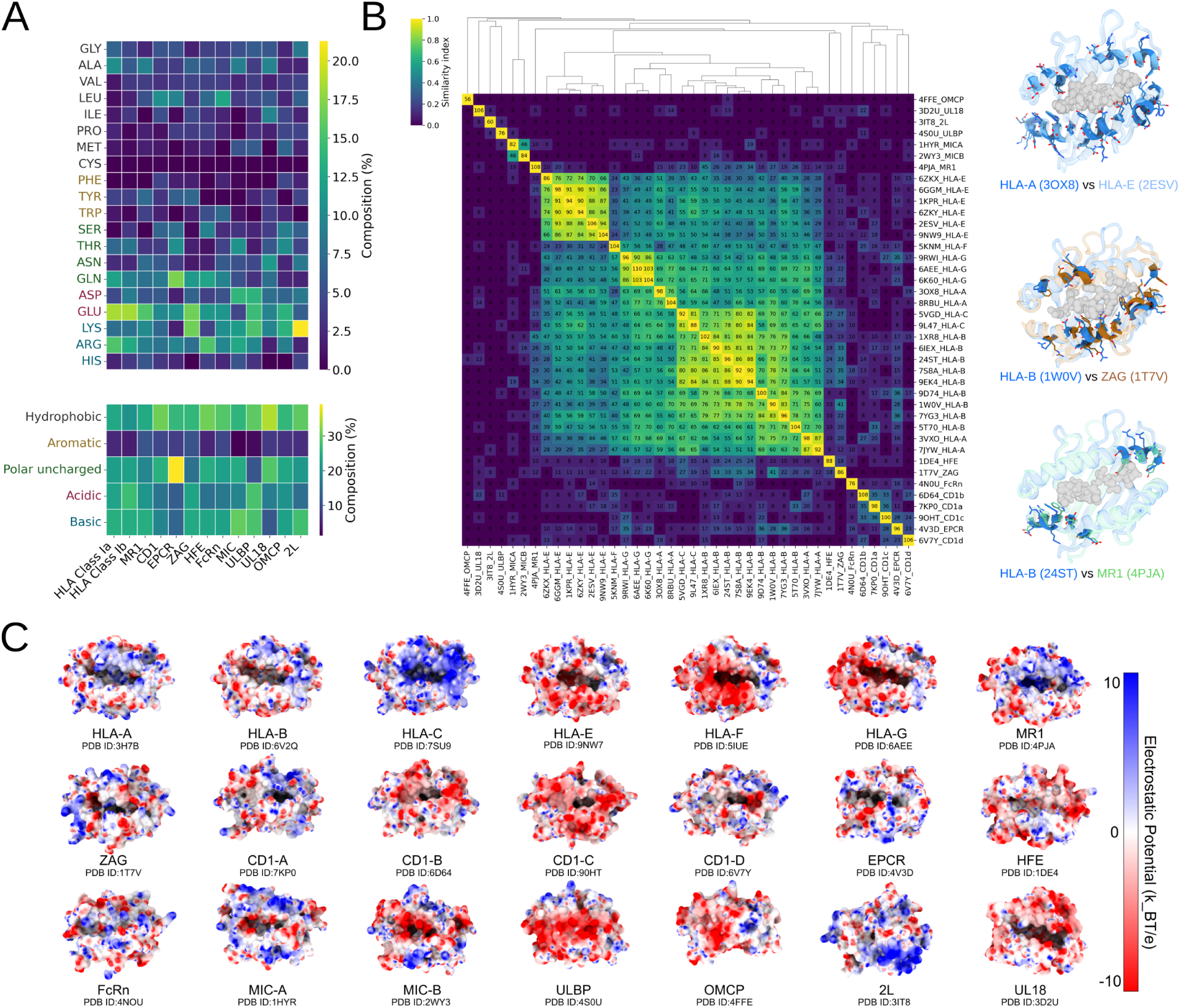
Physicochemical properties of the α1–α2 recognition surface across MHC-fold proteins. (A) Amino-acid composition of solvent-exposed residues within the α1 and α2 helices across MHC-fold protein families. The upper heatmap shows the relative frequency of individual amino acids, whereas the lower heatmap summarizes residues according to physicochemical classes: hydrophobic, aromatic, polar uncharged, acidic, and basic. Frequencies were normalized by the total number of analyzed solvent-exposed helical residues for each family. (B) Pairwise structural similarity among representative MHC-fold protein structures calculated using MHCXGraph. Heatmap colors represent the coverage-based structural similarity index, ranging from 0 to 1, whereas the numerical values within each cell indicate the absolute number of structurally matched residues with identical amino-acid identities. Hierarchical clustering based on the similarity index is shown by the dendrogram. Representative structural superpositions illustrate similarities and differences in the α1–α2 platform between HLA-A and HLA-E, HLA-B and ZAG, and HLA-B and MR1. Bound ligands are shown as spheres where present. (C) Electrostatic surface potential of selected MHC-fold protein structures viewed from the upper α1–α2 recognition surface. Structures were prepared using PDB2PQR, and electrostatic potentials were calculated using APBS and displayed from -10 to +10 kBT/e, with negatively charged regions shown in red, positively charged regions in blue, and approximately neutral regions in white. The PDB ID of each representative structure is indicated below the corresponding model. All quantitative analyses were performed after removal of redundant structures.

Classical MHC-I molecules showed particularly high contributions from Glu and Arg at solvent-exposed positions of the α1–α2 helices (Figure 3A and Supplementary Figure S9). In HLA class Ia, these residues were accompanied by substantial contributions from noncharged residues such as Ala and Gln, resulting in a chemically heterogeneous surface rather than a uniformly charged interface. Differences were also observed within classical MHC-I molecules; for example, HLA-C displayed a higher occurrence of Arg than HLA-A (Supplementary Figure S9A). Non-classical MHC-I molecules showed a modest shift toward acidic character, with HLA-F displaying a particularly negative electrostatic profile in the selected structure shown in Figure 3C.

MHC-like proteins showed substantial variation in the amino-acid composition of their exposed α1–α2 surfaces. Among these proteins, MR1 and ZAG displayed composition profiles most similar to classical MHC-I, clustering adjacent to HLA-A, HLA-B, and HLA-C in the hierarchical analysis of amino-acid composition (Supplementary Figure S9B). Other MHC-like proteins, including CD1, HFE, FcRn, and UL18, showed increased hydrophobic contributions relative to classical MHC. CD1, for example, contained a higher proportion of hydrophobic residues, largely associated with Leu and Ile, whereas UL18 displayed the highest hydrophobic contribution among the analyzed families. Family-specific differences were also evident at finer resolution: CD1a and CD1c displayed distinct electrostatic surface patterns, and MICB exhibited a more pronounced negative electrostatic character than MICA in the selected structures. EPCR showed a particularly distinct composition, with the highest contribution from polar uncharged residues among the analyzed families, driven primarily by Gln and Thr.

Because similar amino-acid composition profiles do not necessarily reflect conservation in the spatial arrangement of residues, we next investigated whether the structural organization of the exposed α1–α2 surface was preserved across MHC-fold proteins. Representative structures were compared using MHCXGraph [52], which identifies geometrically conserved regions containing residues with matching amino-acid identities and quantifies their extent using a coverage-based similarity index (Figure 3B). Classical MHC-I molecules formed a well-defined cluster, indicating substantial conservation in the spatial organization of identical residues within the α1–α2 platform. In contrast, MHC-like proteins generally exhibited lower conservation relative to classical MHC-I and to one another. Nevertheless, conserved regions were identified between selected MHC-like proteins and classical MHC-I molecules, including ZAG, EPCR, and MR1, despite their distinct biological functions and ligand-binding properties.

Together, these analyses demonstrate that conservation of the MHC α1–α2 architecture does not necessarily imply conservation of its exposed surface chemistry. Despite sharing a common structural scaffold, MHC-fold proteins display substantial differences in amino-acid composition, spatial conservation of equivalent residues, and electrostatic surface potential. Notably, similarities in overall composition do not always correspond to equivalent spatial arrangements of residues, highlighting the importance of considering both chemical identity and structural organization when comparing MHC recognition surfaces. This diversity generates distinct molecular environments within a conserved structural framework, providing the basis for investigating how structurally different binding partners engage the α1–α2 platform.

### 2.4 Distinct modes of molecular recognition across the MHC α1–α2 platform

The exposed α1–α2 platform provides an extended surface for macromolecular recognition, but individual binding partners engage only subsets of this surface and do so through distinct structural and physicochemical interfaces. We therefore characterized the architecture, location, size and geometric complementarity of MHC–binder interfaces represented in the PDB.

The exposed α1–α2 surfaces of MHC-fold proteins are recognized by binders employing markedly different secondary-structure elements, although coil regions constitute the predominant interface component in most analyzed complexes (Figure 4A). This pattern is particularly evident for proteins containing Ig-like domains, in which loops connecting β-strands frequently provide the principal MHC-contacting elements. Accordingly, approximately 80% of the interface residues in conventional TCRαβ and antibody-related complexes were assigned to coil regions, largely reflecting the contribution of their complementarity-determining region (CDR) loops. KIR receptors, which also contain Ig-like domains, similarly displayed coil-dominated interfaces, as did CD1–TCR complexes. Loop-mediated recognition was not restricted to Ig-like scaffolds, however, as several structurally unrelated binders, including echovirus capsid proteins, TNFα, designed DARPins, and US2, also contacted MHC surfaces predominantly through coil regions; a small number of CD1–VHH complexes followed the same pattern. In contrast, NKG2-containing receptors and binders such as E3-19K, UL16, VP90, and TAPBP displayed more mixed interfaces, with substantial contributions from both coil and β-strand elements.

**Figure 4:**
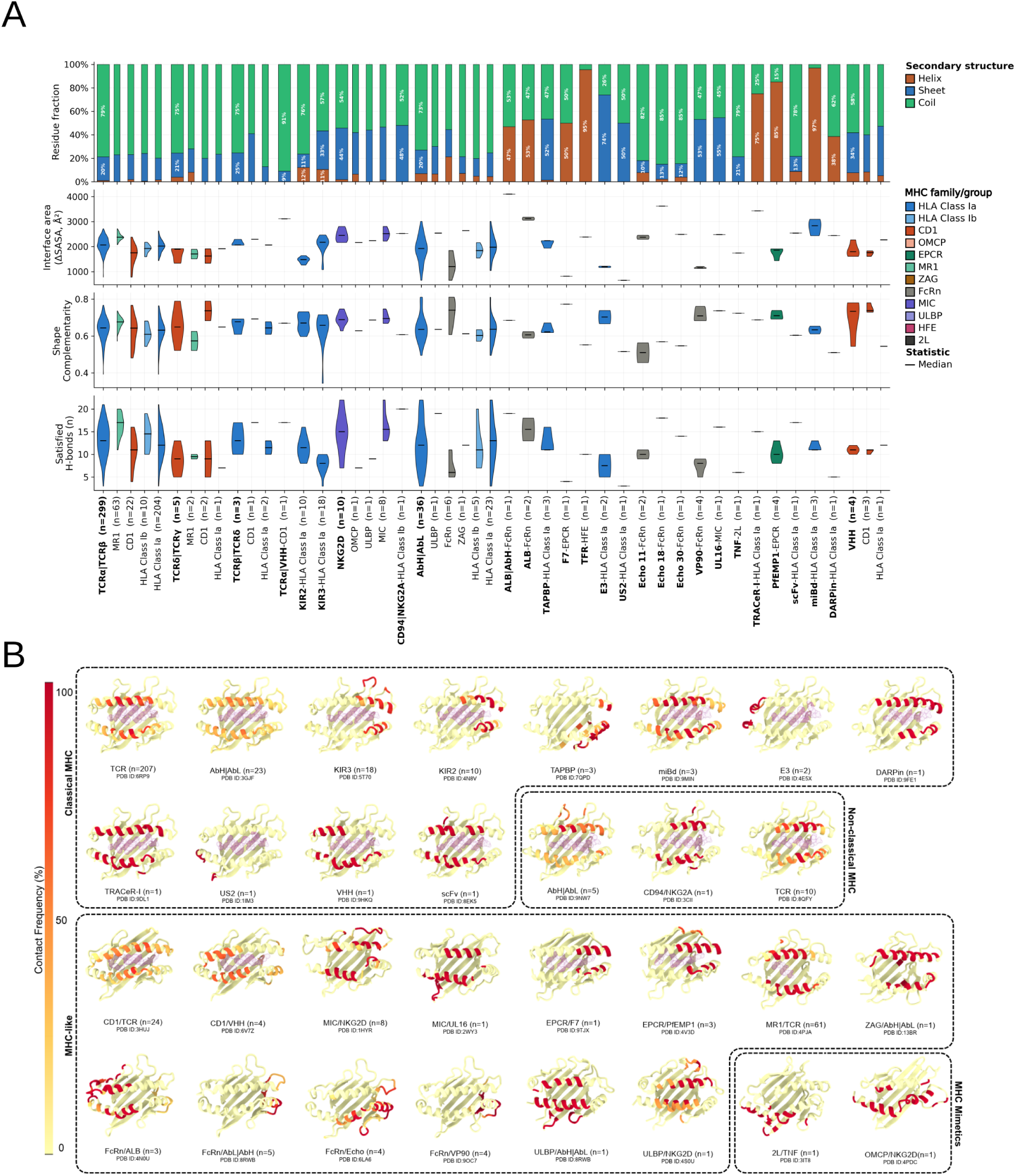
Structural and physicochemical properties of MHC–binder interfaces. (A) Comparison of interface properties across structurally characterized MHC–binder complexes. The upper panel shows the fraction of binder interface residues assigned to α-helix, β-sheet, or coil secondary structure. The subsequent panels show buried solvent-accessible surface area (ΔSASA), shape complementarity, and hydrogen-bond density, expressed as the number of satisfied hydrogen bonds per 1,000 Å² of buried surface area. Binder classes are indicated along the x-axis, with subdivisions according to their interacting MHC families where applicable. Violin plots are colored according to MHC family, and the number of structures analyzed for each group is indicated in parentheses. (B) Spatial distribution of binder contacts across representative MHC α1–α2 surfaces. MHC residues are colored according to the frequency with which they contact the corresponding binder across the analyzed structures, based on residue-level structural correspondence to the group representative. Contact frequencies range from low (yellow) to high (red). The representative structure and the number of complexes contributing to each contact-frequency map are indicated. Bound groove ligands are shown where present. Complexes are organized according to classical ^1^H^2^LA-I, non-classical HLA-I, MHC-like proteins, and viral MHC mimetics. All quantitative analyses were performed after removal of redundant structures.

MHC-fold proteins can also be engaged through markedly different, helix-dominated recognition modes. Natural examples include albumin and PfEMP1-derived binders, whose MHC-contacting interfaces contain substantial α-helical contributions, as well as TfR. A similar strategy has been exploited by engineered binders, including miniproteins and TRACeR-I, which use predominantly α-helical scaffolds to engage the MHC surface (Figures 4A and 5). Together, these observations show that the α1–α2 platform does not impose a single structural solution for molecular recognition and comparable MHC surfaces can be engaged through loop-dominated, mixed coil/β-strand, or α-helix-rich interfaces.

**Figure 5:**
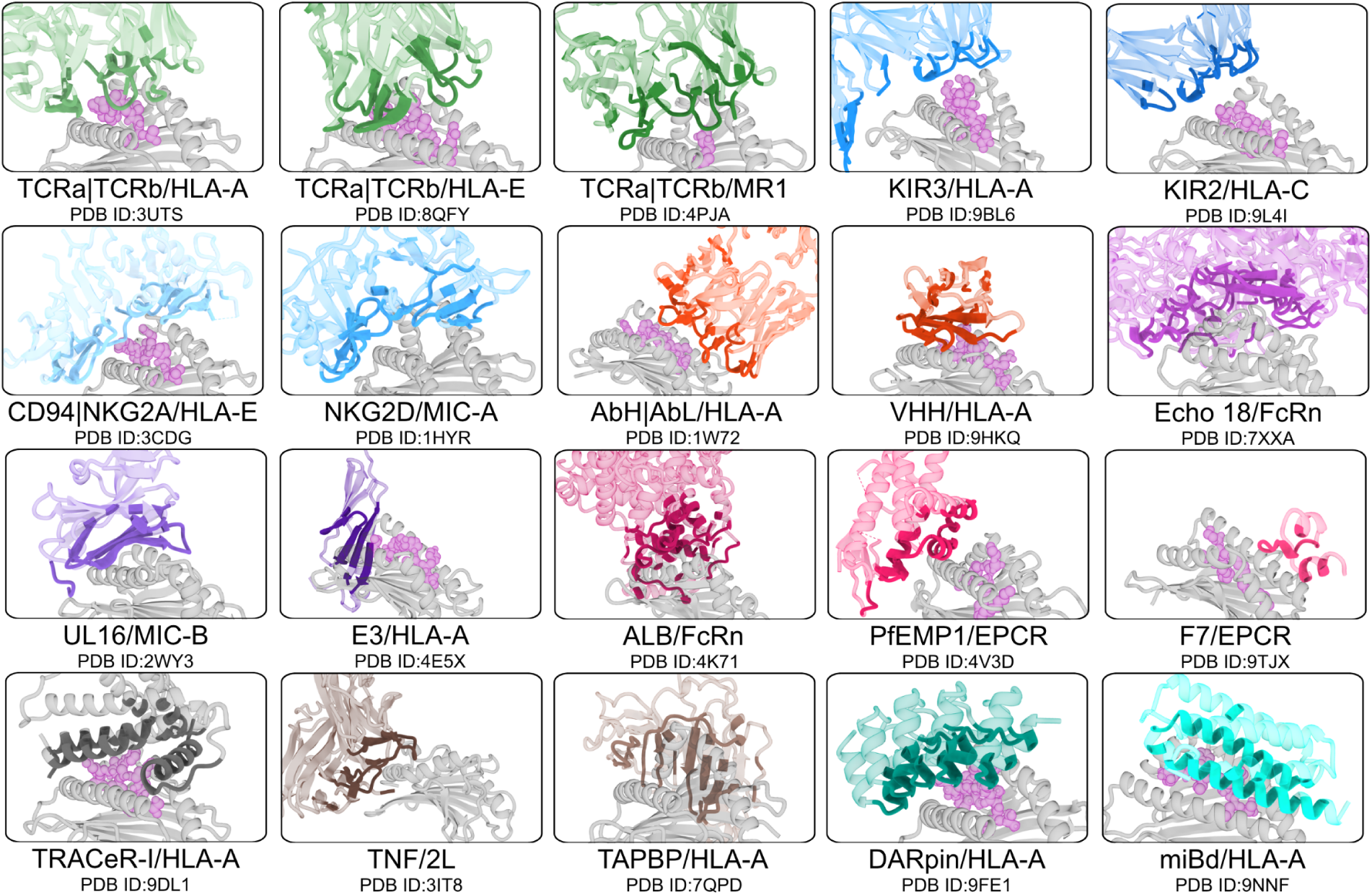
Representative modes of MHC surface recognition by structurally distinct binding partners. Representative MHC–binder complexes illustrate the diversity of recognition geometries observed across the analyzed structural dataset. MHC molecules are shown in gray, with bound groove ligands represented as magenta spheres where present. Binding partners are shown in distinct colors, with MHC-contacting regions highlighted in darker shades. The examples illustrate central *α*1–*α*2 recognition by conventional TCRs, distinct engagement modes by KIR and NKG2-family receptors, antibody-mediated recognition, interactions involving FcRn and EPCR, and recognition mediated by viral proteins and viral MHC mimetics. Engineered MHC binders, including TRACeR-I, DARPin, and miBd, further illustrate alternative structural solutions for engaging the MHC surface. The interacting partners and corresponding PDB identifiers are indicated below each structure.

Beyond differences in binder architecture, the analyzed complexes also revealed distinct spatial modes of engagement across the MHC surface (Figures 4B and 5). Conventional TCR*αβ* complexes predominantly contacted the central regions of both the *α*1 and *α*2 helices, defining the canonical recognition mode of peptide-presenting MHC molecules. TCR recognition of MR1 retained this overall topology but showed a broader footprint, with additional contacts extending toward turns associated with the *β*-sheet platform. Other receptor families also displayed expanded or shifted footprints. Recognition of CD1 by TCRs showed a more heterogeneous spatial distribution of contacts across the α1–α2 platform than observed for classical MHC engagement. KIR3 complexes generally displayed larger interfaces and more extensive footprints than KIR2 complexes, consistent with differences in their overall receptor architectures and modes of MHC engagement. Extended contacts involving regions adjacent to the *β*-sheet floor were also observed for NKG2-family receptors and PfEMP1-derived binders.

A distinct group of binders engaged the MHC predominantly through peripheral regions rather than the central portion of the two helices. These interactions occurred on either side of the *α*1–*α*2 platform and included TAPBP, echovirus capsid proteins, VP90, E3-19K, US2, albumin, TNFα, and the EPCR-binding partners factor VII (F7) and PfEMP1 CIDRα1 domains, although their precise footprints differed substantially (Figures 4B and 5). EPCR provides an interesting example of how structurally distinct binding partners can engage partially overlapping regions of the same molecular surface, as observed for the Gla domain of F7 and the CIDRα1 domain of PfEMP1, despite their different architectures and biological functions. Several viral binders, including US2, E3-19K, echovirus capsid proteins, and VP90, exploit noncanonical peripheral surfaces of the MHC fold rather than reproducing the central recognition geometry characteristic of conventional TCRs. Examples of these recognition modes, highlighting the structural elements contributed by each binder at the interface, are shown in Figure 5.

To further examine the contribution of MHC-bound ligands to recognition, we compared binder–ligand and binder–MHC atomic contacts across complexes containing a resolved ligand (Supplementary Figure S10). Conventional TCRαβ complexes with classical HLA-Ia exhibited a broad distribution of contact ratios, with an average approaching 0.8 and individual values reaching or exceeding 1, indicating substantial direct peptide engagement. However, other

HLA-Ia binders, including KIR2, KIR3, E3-19K, and US2, displayed considerably lower ratios despite recognizing peptide-loaded MHC molecules. Among TCRαβ complexes, direct ligand contacts were generally less prominent for CD1-presented lipids and particularly MR1-presented small-molecule metabolites than for HLA-Ia-bound peptides. For non-classical HLA-Ib, the available CD94/NKG2A complex exhibited a peptide-contact ratio comparable to those observed for TCRαβ. Interestingly, engineered binders, particularly miniproteins (miBd) and DARPin, exhibited relatively high peptide-contact ratios, overlapping with rather than exceeding the distribution observed for conventional TCRαβ complexes. Overall, contacts with the MHC scaffold predominated over direct ligand contacts in most complexes, including those involving engineered binders.

Beyond the differences observed in direct ligand engagement, MHC–binder complexes also exhibited substantial variation in overall interface size, with buried solvent-accessible surface areas (ΔSASA) spanning approximately 600–4,000 Å^2^ (Figure 4A). Conventional TCR*αβ* interactions generally buried approximately 2,000 Å^2^ across the different MHC partners analyzed. Within this group, TCR complexes with MR1 tended to form larger interfaces than those involving classical MHC-I, consistent with the broader footprint observed in the contact-frequency analysis. KIR receptors provided another example of variation within a receptor family, with KIR3 complexes displaying larger interfaces than KIR2 complexes and broader engagement of the MHC surface.

Some of the smallest interfaces were associated with peripheral binding modes, including those formed by the cytomegalovirus protein US2 and the adenovirus E3-19K with classical MHC-I molecules, the interaction between F7 and EPCR, as well as antibodies engaging lateral regions of FcRn and TCRs displaying unusual orientations relative to classical MHC-I (Figure 4A). In contrast, some of the largest interfaces involved FcRn complexes in which albumin and antibody-derived partners contacted an extended surface beyond the upper helical platform. Antibodies displayed one of the broadest distributions of interface areas, consistent with their diverse orientations and functional contexts, which include TCR-mimetic recognition as well as antibodies targeting sites used by natural binding partners such as in FcRn and ULBP. Their structural diversity was reflected in recognition modes ranging from central engagement of the *α*1–*α*2 platform to highly peripheral binding orientations (Figure 4B). More generally, no simple correspondence was evident between the secondary-structure composition of the binder interface and total buried surface area. Helix-rich binders, for example, could generate intermediate-to-large interfaces, as observed for albumin, TfR, miBd, and TRACeR-I, indicating that extensive MHC engagement is not restricted to loop-dominated recognition modes (Supplementary Figure S11A). Consistent with the scope of this study, interface areas were calculated specifically for the MHC *α*1–*α*2 region, although some binding partners may also establish additional contacts with other regions of the complete MHC complex.

Interface size showed little linear association with geometric complementarity across the analyzed complexes (Pearson *r* = 0.03; Supplementary Figure S11B), indicating that, in the analyzed cases, larger buried interfaces were not necessarily associated with higher shape-complementarity values. Most complexes nevertheless displayed shape-complementarity values above 0.6, showing that structurally distinct binder architectures can achieve comparably interfaces with substantial geometric complementarity. Relatively low complementarity was observed for some complexes involving echovirus capsid proteins, US2, DARPins, and TfR, whereas high values were observed in complexes involving F7, TNF, and some antibodies bound to FcRn. Notably, several highly complementary interfaces involved relatively small contact areas, further indicating that interface size and geometric packing represent independent properties of MHC recognition.

To compare polar interactions independently of interface size, we normalized the number of cross-interface hydrogen bonds by buried surface area. Across the complete dataset, interfaces contained on average 6 hydrogen bonds per 1,000 Å^2^ of buried surface, with the central 50% of observations ranging from approximately 4.9 to 7.3 hydrogen bonds per 1,000 Å^2^ (Figure 4A). Conventional TCR*αβ* interfaces were close to this overall distribution, averaging approximately 6.3 hydrogen bonds per 1,000 Å^2^. KIR receptors provided a notable contrast: despite their smaller interface areas, KIR2 complexes displayed a higher hydrogen-bond density than KIR3 complexes. Differences were also observed among complexes involving the same receptor but different MHC-fold proteins. For example, NKG2D interfaces with MIC displayed higher hydrogen-bond densities than the available NKG2D complexes with OMCP or ULBP. Relatively low hydrogen-bond densities were also observed for some engineered binders, including DARPins and designed miniproteins, as well as for the TNF–2L complex.

Together, these analyses reveal a broad diversity of molecular recognition modes across the MHC α1–α2 platform. Structurally distinct binding partners engage different regions of this conserved scaffold through interfaces that vary substantially in secondary-structure composition, footprint distribution, buried surface area, geometric complementarity, and hydrogen-bond density. Notably, comparable regions of the MHC surface can be recognized by binders employing different structural elements, while differences in interface size do not necessarily correspond to differences in geometric complementarity or hydrogen-bond density. These observations highlight the versatility of the α1–α2 platform in accommodating distinct molecular recognition strategies, from canonical TCR engagement to alternative interactions mediated by natural, viral, and engineered binding partners.

## 3 Discussion

Classical MHC-I recognition by T-cell receptors (TCRs) represents one of the most extensively characterized molecular recognition systems involving the MHC fold. The accumulation of experimental data, organized in resources such as VDJdb and TCR3D [53, 54], has supported increasingly detailed analyses of TCR–peptide–MHC recognition and the development of new computational approaches for studying these interactions. More recently, structural knowledge of natural MHC-binding interfaces has also supported protein engineering strategies, including *de novo* designed binders and repurposed protein scaffolds capable of recognizing peptide–MHC complexes [48, 50, 49]. However, classical TCR–pMHC recognition represents only one of the many interaction modes supported by the broader MHC fold.

Despite the extensive knowledge accumulated on classical MHC-I molecules, the broader diversity of proteins containing related α1–α2 domains remains comparatively much less explored, particularly regarding their molecular recognition interfaces and binding partners. Non-classical MHC and MHC-like proteins extend the functions associated with this structural platform to diverse biological processes, ranging from immune regulation and NK-cell-mediated responses to lipid and iron metabolism, while pathogens have also exploited MHC-like architectures and MHC-binding strategies for survival and immune evasion [2]. Motivated by the functional versatility of this scaffold, we performed a systematic structural characterization of MHC proteins from multiple complementary perspectives, encompassing the architecture of their α1–α2 domains, the geometric and physicochemical properties of their ligand-binding cavities and exposed recognition surfaces, and the structural characteristics of their binding interfaces. This work builds on decades of structural studies of individual MHC and MHC-like proteins, taking advantage of the growing availability of experimentally determined structures to provide a broader comparative view of the MHC fold. Importantly, we extend this characterization to the binding partners themselves, systematically examining their structural architectures, recognition footprints, and interface properties across different MHC families. By integrating the MHC scaffold, its recognition surface, and the structural features of its binding partners within a unified framework, our study provides a perspective on how a broadly shared molecular architecture supports diverse modes of recognition across biological systems.

Our structural survey recovered representatives of the currently recognized human classical, non-classical, and MHC-like protein families within the experimentally determined PDB dataset. Classical MHC-I structures included HLA-A, HLA-B, and HLA-C, although their representation was highly uneven across loci and allele groups, reflecting the concentration of available structures in a limited subset of HLA molecules. Among the MHC-like proteins, CD1e was represented by an engineered single-chain construct rather than a native-form complex. Complementary searches of AlphaFold Database models did not identify additional human MHC-fold protein families beyond those already recognized. Although this result suggests that known MHC proteins account for most of the detectable MHC-like structural space within the human proteins surveyed, it should be interpreted within the structural-search criteria limitations of our approach. Pathogen-derived MHC-like proteins constitute an additional source of structural diversity that may be further expanded through searches targeting viral and other microbial proteomes.

Regarding binding partners, most MHC protein families represented in the PDB have been structurally characterized in complex with at least one physiological partner engaging the α1–α2 region considered in our analysis. ZAG and UL18 represent distinct exceptions: no physiological binder engaging the upper α1–α2 surface of ZAG has been structurally characterized, with only an antibody-derived complex represented in our dataset, whereas the known UL18–LILR complexes involve regions outside the recognition surface considered here. Despite the broad repertoire of MHC–binder complexes currently available, several experimentally validated interactions remain without experimentally determined complex structures. Examples include CD94/NKG2C recognition of HLA-E [11], specific γδ TCRs recognizing EPCR [55], TIM-family proteins interacting with phosphatidylserine-presenting CD1d [56], and MICB recognition by NKG2D [28]. Pathogen-derived binding partners further expand this list, including HCMV UL16 binding to ULBP1 and ULBP2 [38], HCMV UL142 interacting with MICA [57, 58], and human herpesvirus 6B (HHV-6B) U20 binding to ULBP1 [59]. Interestingly, both UL142 and U20 have been proposed to adopt MHC-I-like folds [60, 61], suggesting potential examples of interactions between two MHC-like proteins that have not yet been experimentally characterized at atomic resolution. Determining the structures of these unresolved complexes would further expand the current landscape of MHC recognition and provide additional insights into the diversity, organization, and molecular determinants of MHC–binder interfaces.

The α1–α2 platform of MHC proteins integrates multiple structural and physicochemical features that together shape ligand accommodation and binder engagement. Across the analyzed families, this platform spans a continuum of helix openness, from the more widely separated α1 and α2 helices of classical MHC-I and MR1 to more compact arrangements in FcRn, ULBP, and viral MHC mimetics such as OMCP and 2L. Although this geometric variation is broadly associated with differences in ligand-binding capacity, helix separation alone does not determine groove architecture. CD1 and EPCR, for example, combine relatively short helix-to-helix distances with deep cavities and large groove volumes. Conversely, MR1 exhibits a helix separation comparable to classical MHC-I but a smaller average cavity volume, retaining a proportionally larger fraction of that cavity upon binding small-molecule metabolites. These observations indicate that helix geometry, cavity architecture, and ligand occupation represent complementary aspects of the MHC fold, enabling distinct strategies for accommodating chemically diverse molecules. Within this structural landscape, MR1 and ZAG are among the MHC-like proteins most similar overall to classical MHC-I, despite being encoded outside the MHC locus [2]. Both proteins also retain relatively similar amino-acid composition profiles at the exposed α1–α2 surface. However, our analysis of spatial residue conservation revealed differences that are not captured by overall amino-acid composition, highlighting the importance of considering both the chemical properties and spatial organization of recognition surfaces. Interestingly, the structural resemblance of MR1 to classical MHC-I is accompanied by a related mode of TCR engagement, as demonstrated by structural studies of MAIT TCR–MR1 complexes [62, 63]. Nevertheless, geometric and physicochemical similarity of the α1–α2 platform does not necessarily imply equivalent recognition mechanisms or cross-reactivity by the same binding partners. Molecular recognition also depends on the spatial distribution of interacting residues, the identity and chemistry of bound ligands, and additional molecular interactions involving regions outside the upper α1–α2 surface. In classical MHC-I, for example, the CD8 co-receptor engages regions including the α3 domain and β2-microglobulin [64]. Moreover, while classical MHC-I presents peptides and MR1 presents small-molecule metabolites, ZAG can accommodate hydrophobic molecules within its groove [65], although the contribution of these molecules to physiological protein recognition remains unclear. Together, MR1 and ZAG illustrate that substantial functional diversification can occur without major changes in the overall geometry or amino-acid composition of the α1–α2 platform. Conversely, close structural similarity to classical MHC-I is not required for recognition by related receptor families. CD1 molecules display substantial geometric and physicochemical differences from classical MHC-I, yet are recognized by TCRs through diverse interaction modes, highlighting the structural plasticity of TCR-mediated recognition. A complementary example is provided by NKG2D, which recognizes structurally distinct members of the MIC and ULBP families [66, 67], as well as the viral MHC-like protein OMCP [31]. These examples demonstrate that considerable structural and physicochemical variation within the MHC-like scaffold can be accommodated by a common recognition system, emphasizing that molecular recognition cannot be inferred from overall fold similarity alone.

Despite the large exposed surface available for protein recognition, individual binding partners typically engage only a fraction of the α1–α2 platform. Conventional αβ TCRs, for instance, predominantly contact the central regions of both α1 and α2 helices, consistent with the canonical recognition of peptide-presenting MHC-I complexes. This geometry generally positions TCRs near the exposed groove ligands, although their direct contribution to recognition varies across MHC families and individual complexes, and is not always required [68, 69]. In contrast, other binding partners exploit more peripheral regions of the platform, as observed for the adenovirus E3-19K and HCMV US2 proteins [35, 37], whose complete interfaces also extend beyond the α1–α2 domains. A particularly interesting example is HCMV UL16, which promotes the intracellular retention of MICB and selected ULBP proteins while engaging the upper α1–α2 surface of MICB through an interface that substantially overlaps the recognition region used by NKG2D [38]. KIR receptors exhibit intermediate recognition geometries, contacting regions toward the ends of the MHC helices and, in some cases, directly engaging the presented peptide [70]. Within this receptor family, KIR3 complexes generally display broader footprints extending farther toward peripheral regions of the α1–α2 platform than KIR2 complexes, consistent with their distinct domain architectures [71]. Collectively, these interactions demonstrate that structurally and functionally diverse binding partners can recognize a broad range of regions across the exposed α1–α2 surface, extending well beyond the canonical TCR docking region.

Importantly, the α1–α2 domains of MHC proteins are also subject to N-linked glycosylation, as evidenced by multiple experimentally determined structures. Our analysis of structures from the PDB revealed that glycosylation sites are predominantly located along peripheral regions of the α1–α2 platform, although some occur closer to known binder footprints, as observed for CD1d and EPCR. In classical MHC-I, the conserved N-linked glycan is generally positioned outside the canonical TCR-binding footprint, suggesting a limited direct contribution to conventional TCR recognition [72]. Nevertheless, glycosylation has been shown to modulate recognition by other receptors, including KIR3DL1 binding to HLA-B*57:01 [73], as well as T-cell responses involving CD1d [74]. In other MHC-like proteins, such as EPCR, the proximity of glycans to binder footprints suggests potential contributions to molecular recognition, although their precise functional roles remain unclear. Thus, glycosylation represents an additional feature of MHC recognition surfaces that should be considered when interpreting natural interactions and designing new proteins to modulate MHC-associated biological functions.

In most complexes analyzed here, the binder interface with the α1–α2 platform was dominated by residues classified as coil. This pattern was largely driven by binders containing Ig-like domains, particularly TCRs, antibodies, and KIRs, whose exposed loops provide the principal MHC-contacting elements. Outside these structural classes, relatively few examples of similarly loop-dominated recognition of the central upper α1–α2 surface were observed. For classical MHC-I, loop-mediated recognition also appears to be the predominant natural solution for peptide-dependent engagement, particularly among the extensively characterized TCR complexes. In contrast, the available complexes involving non-classical MHC and MHC-like proteins illustrate a broader repertoire of secondary-structure solutions. C-type lectin-like NKG2 receptors and the Ig-like viral immunoevasin UL16, for example, employ substantial β-strand contributions to engage the α-helical platform, whereas α-helical proteins such as albumin recognize distinct surfaces of MHC-like molecules [75]. PfEMP1 provides another interesting example, using a conserved α-helical recognition motif to engage the α1–α2 platform of EPCR [44]. These alternative recognition modes also illustrate that interface architecture is not necessarily associated with its size or geometric packing, as reflected by the lack of correlation between buried surface area and shape complementarity in our dataset.

The diversity of natural recognition architectures also provides inspiration for engineered MHC binders. The unconventional interaction of the *Mycoplasma arthritidis* superantigen MAM with MHC-II inspired the development of the TRACeR platform, subsequently adapted into TRACeR-I, in which an engineered helical scaffold enables peptide-dependent recognition of classical MHC-I complexes [76, 50]. More recently, *de novo* designed α-helical binders targeting classical pMHC-I have demonstrated that compact non-Ig scaffolds can achieve high peptide specificity [49]. Together, these advances highlight how distinct structural solutions can be exploited to engineer MHC recognition, suggesting that similar design principles could be explored beyond classical MHC-I to generate binders capable of engaging and modulating diverse MHC-like proteins.

In conclusion, this study provides an integrated structural landscape of classical, non-classical, and MHC-like proteins, including pathogen-derived MHC proteins, together with the diverse modes by which these molecules are recognized by natural and engineered binding partners. By comparing the architecture, ligand-binding cavities, and physicochemical properties of the α1–α2 platform with the structural characteristics and recognition interfaces of its binding partners, we reveal how a broadly shared MHC fold supports distinct modes of molecular recognition. Our analyses highlight both conserved and divergent structural features across MHC proteins and their binding interfaces, providing a comparative framework for understanding the molecular basis of these interactions. These insights may also help guide the development of novel MHC-targeting binders capable of modulating diverse biological functions, particularly through emerging computational and generative protein-design approaches.

## 4 Methods

### 4.1 Structure-based search for MHC-fold proteins

To identify proteins sharing the characteristic MHC α1–α2 domain architecture, we performed structure-based searches using Foldseek [77], obtained from https://mmseqs.com/foldseek/foldseek-linux-gpu.tar.gz on July 1st 2026. The initial query dataset consisted of 923 human MHC class I structures obtained from TCR3D [54], each containing only the α1-α2 domain. These structures were used as queries to search for structurally related proteins in the Protein Data Bank (PDB) [78] and the AlphaFold Protein Structure Database (AFDB) [79, 80].

For the PDB search, biological assembly files in mmCIF format were retrieved from the RCSB Protein Data Bank (rcsb.org) via its rsync archive (rsync.rcsb.org::ftp_data/assemblies/mmCIF/divided/) on July 30, 2026. A local Foldseek database was constructed from these structures using the createdb command with –chain-name-mode 1, followed by database indexing using createindex with default parameters. For the AFDB search, the prebuilt AlphaFold/UniProt structural database distributed through Foldseek (AFuniprotv6 2025_03), comprising approximately 240 million predicted protein structures, was used. Structural searches against both databases were performed using the easy-search module with –exhaustive-search 1 and –chain-name-mode 1.

### 4.2 Selection of MHC-fold proteins

Structures identified through Foldseek searches were filtered according to the statistical significance and length of their structural alignments to the MHC α1–α2 query structures. Only hits with a Foldseek E-value *≤* 0.01 and an alignment length (alnlen) *≥* 90 residues (corresponding to half of the α1–α2 domain) were retained for the subsequent analyses. The alignment-length threshold was applied to reduce the inclusion of short or partial structural matches, including those involving individual MHC-II chains, while prioritizing matches covering a substantial portion of the MHC α1–α2 architecture. To evaluate the suitability of the selected E-value threshold, we additionally examined its relationship with query-normalized TM-scores (qTM-scores), using a value of 0.5 as a reference for structural fold homology. This comparison indicated that the selected E-value cutoff was broadly consistent with the identification of structurally similar proteins (Supplementary Figure S12). Importantly, TM-scores were used only for comparative evaluation and were not applied as an additional filtering criterion.

### 4.3 Processing and standardization of MHC-fold structures

The PDB or UniProt identifiers (the latter for AFDB entries) corresponding to the selected hits were used to retrieve the respective structures in mmCIF format. For PDB entries, the first biological assembly was selected from the Foldseek-filtered hits, as its availability was confirmed across all PDB entries. To identify potential inconsistencies in biological assembly composition that could result in the omission of MHC-associated peptides or macromolecular binding partners, all biological assemblies reported for each PDB entry were retrieved based on the RCSB Data API’s assembly_count field. The polymeric entity composition of each alternative assembly was compared with that of the selected assembly using the _entity and _struct_asym mmCIF categories. Entries containing polymeric entities in alternative assemblies that were absent from the selected assembly were flagged for manual inspection and excluded when assembly inconsistencies were confirmed. Assemblies differing only in the number of copies of an otherwise identical set of polymeric entities were not flagged.

Structures containing extensive unresolved regions, incomplete or incorrectly assigned biological assemblies, binders from species outside the scope of the study, or structural and coordinate quality issues were excluded following manual inspection (Supplementary Table 1). Structures containing post-translational modifications that interfered with downstream analyses were also excluded during this curation step. N-linked glycans were evaluated separately prior to the removal of non-protein heteroatoms, as described in Section 4.11.

For PDB entries containing multiple copies of the selected MHC-fold protein within the retained biological assembly, a single representative chain was selected. The first-ranked chain in the E-value-sorted Foldseek alignment file was designated as the primary MHC chain, while the remaining copies were treated as duplicates and excluded from subsequent processing and analysis. To generate the MHC-only representation, all other chains were removed, and the primary MHC chain was renumbered to match the 1-indexed target positions reported by Foldseek. Residues outside the alignment interval defined by tstart and tend were subsequently discarded. This procedure, applied to both the PDB and AFDB datasets, delineated the continuous structural region corresponding to the MHC α1–α2 platform in each selected structure, providing a consistent structural representation for downstream comparative analyses.

Each PDB MHC-only structure was aligned to a reference MHC-I structure (PDB ID: 1AO7) using TM-align [81], yielding a rigid-body transformation that placed it in a common, consistently oriented reference frame. In parallel, the selected biological assembly was subjected to an initial spatial filtering step using Biopython’s NeighborSearch module. Each chain other than the primary MHC chain was retained only if at least one of its atoms was within 10 Å of any Cα atom of the primary MHC chain. This procedure served as a preliminary selection of potential ligands and macromolecular binding partners, whose identification and annotation were performed in subsequent processing steps. The retained chains were renamed for consistency, with the primary MHC chain designated as chain A. The rigid-body transformation obtained from the MHC-only alignment was subsequently applied to the filtered assembly, placing it in the same reference frame as the MHC-only representation. The aligned MHC-only structures were used for cavity detection, as described in Section 4.13.

Structural processing of AFDB models was considerably simpler than that of PDB entries, as these models represent individual proteins without associated ligands or macromolecular binding partners and do not require reconstruction of unresolved coordinate regions. Accordingly, AFDB structural processing was limited to the extraction and standardization of the MHC *α*_1_–*α*_2_ domain, whereas biological assembly validation, ligand and binder identification, and reconstruction of unresolved structural regions were performed exclusively for PDB structures. The resulting AFDB MHC-only representations were retained for subsequent dataset filtering, functional annotation, and structural similarity comparisons.

### 4.4 Identification of MHC ligands and binding partners

Following structural processing, ligands and macromolecular binding partners associated with the selected MHC-fold proteins were identified in structures obtained from the PDB. This procedure was also used to annotate the ligand-bound and binder-associated states of the analyzed MHC structures.

Groove-bound ligands, including peptides, lipids, and other molecules, were identified based on their spatial proximity to cavities detected using pyKVFinder [82, 83, 84], following the cavity detection procedure described in Section 4.13. For each structure, cavities were identified using the corresponding MHC-only representation. Residues in the filtered assembly located within 1 Å of any point of the detected cavities were considered candidates for ligand assignment. MHC residues outside the α1–α2 domain and residues from other protein chains that exhibited contacts with the cavity were further investigated to identify residues potentially misassigned to these chains. Non-amino-acid residues satisfying this criterion were automatically reassigned to a separate chain. In turn, amino-acid residues were considered for reassignment only when cavity contacts involved backbone atoms and were reassigned following manual confirmation. Contacts involving exclusively side-chain atoms were not considered sufficient evidence of incorrect chain assignment. Water molecules, crystallization artifacts, and other non-biological components (Supplementary Table 2) were removed from the selected structures.

Following ligand identification and chain reassignment, the primary MHC chain in the filtered assembly was restricted to the α1–α2 domain using the same Foldseek-derived alignment positions (tstart and tend) employed to generate the MHC-only representation (Section 4.3).

The macromolecular binding partners were subsequently identified by simultaneously meeting two criteria: (1) a minimum chain length of 30 residues and (2) a distance of 4 Å or less to helical residues of the MHC chain or to a previously identified groove-bound ligand. The reference set of MHC helical residues was obtained using DSSP, considering all residues within the retained α1–α2 domain assigned to helical secondary-structure classes (H, G, or I). Chains satisfying these criteria were retained together with the MHC chain and the confirmed groove-bound ligands, while all other chains were removed.

The presence of multiple copies of an MHC-containing complex within the same biological assembly may lead to incorrect binder assignment during contact-based identification, as a binding partner associated with one MHC copy may also establish spatial contacts with another. To identify potentially ambiguous cases, two complementary criteria were employed. First, the global stoichiometry of each selected biological assembly was retrieved from the RCSB GraphQL Data API (rcsb_struct_symmetry). Assemblies in which all distinct polymer chain types occurred at the same multiplicity greater than one (e.g., A2B2C2D2) were flagged as potentially containing multiple copies of the same complex. Second, entries containing multiple MHC-fold chains identified by Foldseek were additionally flagged, including those that did not satisfy the stoichiometric criterion. These complementary criteria were used to identify cases requiring manual inspection of the spatial organization of MHC chains and their associated binding partners. When a chain was confirmed to be a binding partner of another MHC copy (e.g., copies of the TCR or β2-microglobulin), only that chain was removed, and the structure was retained in the dataset (Supplementary Table 3).

### 4.5 Identification and modeling of unresolved structural regions

Unresolved regions in the selected PDB structures were identified by combining sequence alignment with an evaluation of backbone continuity. Reference sequences were extracted from the SEQRES records of the original PDB files or the corresponding polymer entity information in mmCIF files. Geometric backbone breaks were identified when the distance between the carbonyl carbon and the backbone nitrogen of consecutive observed residues exceeded 2.5 Å. The observed and reference sequences were globally aligned using Biopython’s PairwiseAligner (version 1.87) [85], with gap opening, extension, and terminal gap scores of *−*12, *−*10, and *−*10, respectively. When multiple equally optimal alignments were obtained, a custom heuristic scoring function was employed to select the alignment based on gap distribution, alignment density, continuity of observed segments, and consistency with geometric backbone breaks. Ambiguous cases were manually inspected and curated. This procedure was not applied to AFDB models, which are predicted as complete chains and therefore do not contain unresolved regions.

Unresolved internal regions were reconstructed with Rosetta (version 2024.09) [86] through a two-step modeling protocol, using the ref2015 score function [87]. Missing residues and their flanking anchor residues were mapped onto a Rosetta blueprint, with amino acid identities specified using the PIKAA directive. N-terminal and C-terminal gaps were not modeled. Initial reconstruction was performed using the Rosetta Remodel application with the low-resolution quick_and_dirty protocol. Between 50 and 1,000 independent trajectories were generated per target, with 50 trajectories used as the standard procedure. The five lowest-energy models were retained, and the lowest-energy model was selected for subsequent refinement.

The selected model was refined in full-atom mode using the Rosetta LoopModel application with the kinematic closure (KIC) refinement protocol (refine_kic). Refinement encompassed the reconstructed regions and their flanking anchor residues, with enhanced rotamer sampling enabled through the ex1 and ex2 options. Side-chain conformations of neighboring residues were preserved using the fix_natsc option.

### 4.6 MHC-fold dataset filtering and structural grouping

The resulting dataset of MHC-fold protein structures and complexes was filtered to retain proteins from *Homo sapiens* and pathogen-derived MHC-fold proteins associated with human infections and interactions with human proteins. The selected proteins were classified according to their corresponding protein families and broader functional and structural categories, as described in Section 4.7.

Additional filtering criteria were applied to structures obtained from the AFDB. Models associated with UniProt identifiers flagged for exclusion from the UniProt database were removed [88]. Only models with aligned target regions corresponding to the α1–α2 platform exceeding 140 residues in length were retained. This threshold was established based on the sequence length distribution observed among the PDB hits. Furthermore, structural comparisons between AFDB models and experimentally determined PDB structures were restricted to reviewed Swiss-Prot entries to ensure greater reliability of the associated protein annotations.

For downstream analyses, the processed structures were organized into four structural representations: MHC-only, MHC–ligand, binder-only, and MHC–ligand–binder. Only MHC-only representations were generated for AFDB models, which were used for structural similarity comparisons with experimentally determined PDB structures and for the assessment of model confidence. Subsequent quantitative analyses of MHC structural and physicochemical properties, ligand-binding cavities, and MHC–binder recognition interfaces were restricted to experimentally determined PDB structures, with redundancy reduction applied where indicated.

### 4.7 Protein identification and functional annotation

Identification and functional annotation of the selected MHC-fold proteins and their binding partners were performed using information retrieved from publicly available biological databases, complemented by manual curation. For PDB entries, the Uniprot identifier corresponding to each chain was obtained using the Structure Integration with Function, Taxonomy and Sequences (SIFTS) resource [89]; for AFDB models, the UniProt identifier was obtained directly from the model identifier. In both cases, protein names, source organisms, gene names, and UniProt entry status were subsequently retrieved for the corresponding UniProt identifier through the UniProt REST API [88] (queried August 19, 2026). For binding-partner chains (Section 4.4), descriptions that remained incomplete or ambiguous were supplemented using polymer-entity descriptions retrieved from the RCSB Protein Data Bank GraphQL Data API [90].

Protein identities were established primarily based on gene annotations. The gene names retrieved from UniProt were standardized and assigned to MHC families/groups and categories using a manually curated mapping table. For PDB entries, this was followed by manual curation to resolve inconsistencies in protein nomenclature and supplement missing information when necessary. For AFDB models, protein identities were assigned using a manually curated mapping table. MHC-fold proteins and their binding partners were subsequently organized according to their corresponding protein families and functional groups. Structures for which a specific MHC gene could not be assigned were excluded from the dataset (142 in total for AFDB and 1 for PDB).

PDB structures corresponding to single-chain fusion or chimeric constructs were identified through multiple protein mappings associated with the same structural chain and annotated separately according to their construct type.

Additional structural information was collected to characterize the quality of the analyzed models. For AFDB structures, mean predicted Local Distance Difference Test (pLDDT) values were calculated considering only the extracted MHC α1 and α2 domains. For experimentally determined structures, resolution values were obtained from the rcsb_entry_info.resolution_combined field of RCSB Protein Data Bank API [91].

### 4.8 Identification and distribution of HLA allele groups

To characterize the HLA alleles represented in the structural dataset, PDB entries annotated as HLA-A, HLA-B, HLA-C, HLA-E, HLA-F, or HLA-G were analyzed to identify their corresponding allele groups at the first-field level (e.g., HLA-A*02). For each entry, the *α*_1_–*α*_2_ domain sequence of the MHC chain (chain A) was extracted using Biopython and compared against the IPD-IMGT/HLA protein reference database (hla_prot.fasta, release 3.66.0, October 14, 2026 (*github.com/ANHIG/IMGTHLA*))), considering all reference alleles of the corresponding HLA gene. Allele groups were preferentially assigned based on exact sequence substring matches. When no exact match was found, the closest reference alleles were identified through local pairwise sequence alignment using Biopython’s PairwiseAligner, with the BLOSUM62 substitution matrix, a gap-opening penalty of *−*10, and a gap-extension penalty of *−*0.5. Allele groups were then assigned based on the reference alleles yielding the highest alignment score. As an independent check, the allele reported in the PDB entry title, when available, was compared with the sequence-based assignment. The distribution of allele groups is presented for HLA-A, HLA-B, and HLA-C, the only HLA loci represented by more than one allele group in the dataset. Relative frequencies were calculated considering only structures unambiguously assigned to a single allele group within each locus (Supplementary Figure S2).

### 4.9 Structural similarity analysis of MHC-fold proteins and their binding partners

Pairwise structural comparisons were performed using TM-align [81] with default parameters. Structural similarity was quantified using pairwise average TM-scores, which were organized into symmetric similarity matrices. Structural distances were subsequently calculated as *d_ij_* = 1 *−* TM*_ij_*.

To visualize structural relationships among MHC-fold proteins and, independently, among their macromolecular binding partners, Uniform Manifold Approximation and Projection (UMAP) [92] was applied directly to the corresponding pre-computed structural distance matrices (metric = precomputed). Both two-dimensional embeddings were generated using the same parameters: n_neighbors = 8, min_dist = 0.8, spread = 1.0, repulsion_strength = 1.0, and random_state = 42.

For MHC-fold proteins, a single embedding was generated using the combined dataset of experimentally determined PDB structures and selected reviewed AFDB models, including single-chain fusion constructs and chimeric structures (Supplementary Figure S1A). For the main representation (Figure 1A), AFDB models, single-chain constructs, and chimeric structures were omitted exclusively from the visualization, without recalculating the UMAP or modifying the coordinates of the remaining structures. The structural landscape of MHC-binding partners was generated using the binder-only structures obtained from the PDB (Figure 1C and Supplementary Figure S1B). Protein family and functional annotations were used to characterize structural clustering patterns within both embeddings.

For each MHC family, or individual protein group in the case of classical and non-classical HLA-I, CD1, and MIC proteins, a representative structure was selected as the medoid, defined as the eligible structure exhibiting the highest mean pairwise TM-score relative to all other members of the corresponding group. AFDB models, single-chain fusion constructs, and chimeric structures were excluded as representative candidates but remained included in the pairwise comparisons used to calculate mean structural similarities. Groups containing a single eligible PDB structure were represented by that structure itself.

Structural relationships among the selected representatives were further characterized using pairwise TM-scores and root-mean-square deviations (RMSDs) calculated over the residue positions structurally aligned by TM-align (Supplementary Figure S4). Additional comparisons were performed between reviewed AFDB models and their best-matching experimentally determined PDB structures, between single-chain fusion constructs and their best-matching native structures, and between naturally ligand-free structures and their best-matching ligand-bound counterparts within the corresponding MHC families. These analyses were used to assess structural conservation of the *α*_1_–*α*_2_ platform across different structural sources, construct types, and ligand-binding states (Supplementary Figure S5).

### 4.10 MHC complex redundancy analysis

Redundant PDB structures and complexes were removed from analyses in which the overrepresentation of equivalent complexes could introduce bias. Redundancy was evaluated using the final processed structures, considering the trimmed MHC *α*_1_–*α*_2_ sequence and the associated binding partners and ligands retained during the preceding processing steps. Two complexes were considered redundant only when they satisfied all of the following criteria: (1) their trimmed MHC sequences were identical or one sequence was entirely contained within the other; (2) they contained the same number of binder chains, with a one-to-one correspondence between identical or contained binder sequences; and (3) they exhibited identical ligand compositions. Sequence containment required an exact contiguous match, without substitutions, insertions, or deletions. Peptide ligand sequences and non-peptide ligand Chemical Component Dictionary (CCD) identifiers were compared as multisets, preserving the number of occurrences while disregarding chain order and chain identifiers. Thus, complexes containing the same MHC protein but differing in their binding partners or ligands were retained as distinct entries.

Structures were organized into redundancy groups, in which each member was required to be directly redundant with the designated group reference structure. The selection of reference structures prioritized previously curated structural representatives selected based on TM-score [81]. In the absence of such representatives, structures were prioritized according to (1) the greatest trimmed MHC sequence length, (2) the highest experimental resolution, and (3) the greatest combined amino acid sequence length across all binder chains. Remaining ties were resolved by selecting the structure with the lexicographically smallest PDB identifier. When two previously curated structural representatives were directly redundant, both were retained as reference structures of distinct redundancy groups. Of the 1,267 selected and processed MHC structures from the PDB, 95 were identified as redundant, yielding a reduced dataset of 1,172 structures. This dataset was used in analyses for which redundancy removal was explicitly indicated.

### 4.11 Identification and characterization of N-linked glycosylation

N-linked glycosylation sites were identified directly from experimentally resolved glycan residues in PDB structures, prior to the removal of non-protein heteroatoms during structural processing (Section 4.3). Glycan components were identified based on the presence of the monosaccharide residue types NAG, NDG, MAN, BMA, GLC, BGC, and FUC. An N-linked glycosylation site was identified when any atom of a glycan residue was located within 2.0 Å of the side-chain ND2 atom of an asparagine (ASN) residue.

To avoid counting multiple monosaccharides belonging to the same glycan as independent glycosylation sites, only the root sugar directly attached to the asparagine residue was considered for site identification. The composition of each glycan was characterized using all monosaccharide residues assigned to the same glycan chain. Thus, individual glycosylation instances corresponded to glycan attachment sites rather than individual monosaccharide residues.

Glycosylation sites associated with the primary MHC chain (chain A) were subsequently mapped onto the corresponding trimmed MHC-only structure through pairwise sequence alignment, and sites outside the retained *α*_1_–*α*_2_ domain were excluded. Based on the alignment-derived helix boundaries defined in Section 4.12, glycosylation sites were classified as located within the *α*_1_ helix, the *α*_2_ helix, or outside the two helical regions.

To compare glycosylation sites across structures associated with the same gene, individual sites were mapped through TM-align structural alignments onto a gene-specific reference structure, selected as the glycosylated structure containing the largest number of identified sites. The relative frequency of glycosylation at each mapped position was calculated as the number of structures containing an experimentally resolved glycan at that position divided by the total number of structures associated with the corresponding gene. The spatial distribution of N-linked glycosylation sites and the diversity of their glycan compositions across MHC-fold proteins are illustrated in Supplementary Figure S3.

### 4.12 Characterization of the MHC α1–α2 helical architecture and surface composition

To characterize the architecture of the *α*_1_–*α*_2_ platform across MHC-fold proteins, the two canonical groove helices were mapped using the representative HLA-A structure (PDB ID: 3H7B) as a reference, with residues 51–85 and 136–180 defining the *α*_1_ and *α*_2_ helical regions, respectively. Each PDB MHC *α*_1_–*α*_2_ structure was aligned to the reference using TM-align to establish residue-level structural correspondences. For each structure, the boundaries of each helical region were defined by the minimum and maximum residue positions aligned to the corresponding reference region. The lengths of these regions and their mean nearest-neighbor C*α* distance were subsequently calculated. The latter was defined as the average distance from each C*α* atom in the *α*_1_ region to its closest C*α* atom in the *α*_2_ region and was used to characterize the relative openness of the MHC platform.

To investigate the physicochemical composition of the exposed *α*_1_–*α*_2_ surface, relative solvent accessibility (RSA) was calculated for each MHC residue using DSSP (mkdssp, v4.5.7) [93]. These calculations were performed on structures retaining any groove-bound ligands but excluding macromolecular binding partners. Residues within the mapped *α*_1_ and *α*_2_ helical regions with RSA *>* 0.2 were classified as solvent-exposed. Amino acid composition was determined by pooling solvent-exposed residues across all structures within each group. The relative frequencies of individual amino acids and their corresponding physicochemical classes were calculated by normalizing their counts by the total number of solvent-exposed residues within each group. Amino acids were grouped into five physicochemical classes based on the IMGT chemical classes [94], with sulfur-containing residues merged into the aliphatic group and hydroxyl and amide residues merged into a single polar group: hydrophobic (Gly, Ala, Val, Leu, Ile, Pro, Met, Cys), aromatic (Phe, Tyr, Trp), polar uncharged (Ser, Thr, Asn, Gln), acidic (Asp, Glu), and basic (Lys, Arg, His). Amino-acid composition profiles were standardized per amino acid (z-score) across families, pairwise Euclidean distances were computed, and families were clustered hierarchically using average linkage.

The helical content of each MHC *α*_1_–*α*_2_ domain was calculated independently of the alignment-based helix boundaries using DSSP secondary-structure assignments. Residues assigned to the H, G, and I categories were considered helical, and their proportion relative to the total number of residues within the *α*_1_–*α*_2_ domain was calculated.

### 4.13 Detection and characterization of ligand-binding cavities in MHC-fold proteins

Ligand-binding cavities within the MHC *α*_1_–*α*_2_ platform were identified and characterized using the structurally aligned MHC-only representations. Hydrogen atoms were removed from each structure prior to cavity detection. Cavities were identified using pyKVFinder (v0.9.0) [82], with a probe-in radius of 1.4 Å a probe-out radius of 6.0 Å and a minimum cavity volume cutoff of 30 Å^3^.

Because pyKVFinder identifies cavities throughout a protein structure, a spatial filtering procedure was applied to retain only cavities associated with the region corresponding to the canonical MHC ligand-binding groove. The C*α* atoms of the peptide extracted from the reference MHC-I structure (PDB ID: 1AO7) were used to define this region within the common structural reference frame established during structural processing. A cavity was retained when at least one of its points was located within 1.0 Å of any C*α* atom of the reference peptide. For each retained cavity, pyKVFinder was used to calculate its volume (Å^3^), surface area (Å^2^), maximum depth, and mean depth. When multiple cavities satisfied the spatial filtering criterion within the same structure, their individual volumes were summed to obtain the total cavity volume.

To characterize the extent of cavity occupation by bound ligands, the same cavity detection and spatial filtering procedures were additionally applied to the corresponding MHC–ligand representations, which retained groove-bound ligands but excluded macromolecular binding partners. For each ligand-bound structure, a residual cavity ratio was calculated as the cavity volume detected in the MHC–ligand representation divided by that detected in the corresponding MHC-only representation (*V*_MHC*−*ligand_*/V*_MHC*−*only_). Structures for which no residual cavity was detected in the presence of the ligand were assigned an undefined ratio rather than a value of zero and were excluded from the residual cavity ratio analysis.

### 4.14 Structural conservation analysis of MHC-fold proteins using MHCXGraph

Structural conservation among representative MHC-fold proteins was additionally characterized using MHCXGraph [95] (version 0.1.3), which identifies geometrically conserved regions containing residues with matching amino-acid identities independently of conventional sequence alignment. Pairwise comparisons were restricted to residues within the mapped *α*_1_ and *α*_2_ helical regions exhibiting relative solvent accessibility (RSA) *≥* 0.1. Graphs were constructed using Cα-only node granularity and a distance threshold of 10 Å excluding bound ligands.

For each structural pair, the extent of conservation was quantified using MHCXGraph’s coverage-based similarity index, while the absolute number of structurally matched residues with identical amino-acid identities was also recorded. The resulting pairwise similarity matrix was converted into a distance matrix and subjected to hierarchical clustering using the average-linkage method.

### 4.15 Electrostatic surface potential analysis of MHC-fold proteins

Electrostatic surface potentials were computed for one representative structure per analysis group, selected by the highest average pairwise TM-score relative to the remaining group members [81]. Calculations were restricted to the MHC heavy chain, with peptide, lipid and other groove-bound ligands removed beforehand, so that the reported potential corresponds to the empty groove. Protonation states and atomic parameters were assigned with PDB2PQR (version 3.6.1) [96] using the PARSE force field [97], with titration states predicted at pH 7.4 by PROPKA [98] (version 3.5.1), and missing heavy atoms and hydrogens added by the same program.

The linearized Poisson-Boltzmann equation was solved with APBS (version 1.5) [99], using the default grid parameters of the APBS input generator, with relative dielectric constants of 2 for the protein interior and 78.54 for the solvent, 298 K, a monovalent ionic strength of 150 mM NaCl, a harmonically smoothed solvent-excluded dielectric boundary, and cubic B-spline charge discretization. A sequential focusing scheme was applied, in which a coarse grid spanning 1.7 times the molecular bounding box provided the boundary conditions for a fine grid targeting approximately 0.5 Å per grid point, both derived from the bounding box of each individual structure.

Molecular surfaces were coloured by the resulting potential in UCSF ChimeraX (version 1.11) [100, 101], using a symmetric red-white-blue scale spanning *−*10 to +10 *k*_B_*T/e* that was kept identical across all structures and encompassed the full range of surface potentials observed (Figure 3C).

### 4.16 Structural and physicochemical characterization of MHC-binder interfaces

The structural and physicochemical properties of interfaces between MHC-fold proteins and their macromolecular binding partners were characterized using PyRosetta (version 2026.05) [102]. Prior to interface calculations, each complex was subjected to a single round of all-atom relaxation using the FastRelax protocol [103], with five standard repeat cycles and PyRosetta’s default full-atom score function. All subsequent interface calculations for each complex were performed using the resulting relaxed conformation.

Interface properties were calculated using the InterfaceAnalyzerMover [104], configured with an explicit chain-string specification defining the two interacting sides, the binder and the MHC chain together with its groove-bound peptide or lipid ligand. The mover was configured to repack side chains in the unbound state prior to interface energy recalculation, without repacking the input pose, and to calculate shape complementarity [105] and hydrogen-bond-related solvent-accessible surface area (SASA) properties. The extracted descriptors included interface binding energy (ΔG), buried solvent-accessible surface area (ΔSASA), the side-chain contribution to buried surface area, the number of interface hydrogen bonds, the fraction of unsatisfied buried polar groups, and total and cross-term interface energies. For protein-only interfaces, additional descriptors included packing quality (packstat) [106] and contact molecular surface (CMS) area. Hydrogen-bond density was calculated as the number of satisfied cross-interface hydrogen bonds normalized by buried surface area, expressed per 1,000 Å^2^ of ΔSASA.

Because macromolecular binding partners were identified by a distance criterion relative to the MHC chain and its groove-bound ligand (Section 4.4), individual chains of obligate multi-chain binders occasionally fell outside the contact cutoff despite belonging to the same binding unit, as observed for one chain of some heterodimeric binders. These complexes were analyzed together with those retaining all binder chains.

Interfaces for which the InterfaceAnalyzerMover simultaneously returned zero ΔG, zero ΔSASA, and zero interface residues were considered failed interface detections and excluded from subsequent analyses. Buried surface area, shape complementarity and hydrogen-bond density were compared across binder classes and their corresponding MHC partners (Figure 4A), and the relationship between interface size and geometric complementarity was assessed by Pearson correlation (Supplementary Figure S11B). All interface descriptors refer exclusively to the MHC *α*_1_-*α*_2_ region, so that contacts established by a binder with other domains of the complete MHC assembly are not included.

Binder engagement of the groove-bound ligand was quantified. Engagement was expressed as a contact ratio, defined as the number of binder-ligand heavy-atom pairs divided by the number of binder-MHC heavy-atom pairs, a pair being counted whenever two heavy atoms from opposite entities were located within 4 Å of each other (Supplementary Figure 10). Hydrogen atoms were excluded, as they are absent from most deposited models. Complexes in which a ligand was present but not contacted by the binder were retained with a ratio of zero.

Ligand solvent accessibility was computed with the Shrake-Rupley algorithm [107] as implemented in Biopython [108], using a probe radius of 1.4 Å, Bondi van der Waals radii [109] and 3,000 test points per atom, the latter increased from the default of 100 after verifying convergence of the computed ligand SASA. A ligand heavy atom was considered accessible when its SASA was at least 1 Å^2^. Accessibility was computed on the MHC-ligand assembly alone, thereby reporting the intrinsic exposure of the ligand within the groove, and reported as the absolute number of accessible ligand heavy atoms summed over all ligand chains (Supplementary Figure S10B), with the corresponding percentage of accessible atoms retained as an alternative representation; the equivalent calculation including the binder, which instead quantifies ligand occlusion, was computed for comparison. Distributions of both descriptors were displayed as kernel density estimates grouped by binder class and subdivided by MHC class, annotated with medians and quartiles, using Matplotlib [110].

### 4.17 Spatial distribution of binder contacts across representative MHC α1–α2 surfaces

Physical contacts between the MHC *α*_1_-*α*_2_ region and each macromolecular binding partner were identified directly from atomic coordinates using Biopython’s NeighborSearch module [111]. A residue was considered part of the interface when at least two of its atoms were located within 4 Å of any atom of the opposing interaction partner. Secondary-structure assignments for binder interface residues were obtained using pydssp [93, 112] and used to characterize the relative contributions of helical, *β*-strand, and coil regions to the binding interface (Figure 4A). Based on these assignments, interfaces were classified according to the predominant secondary-structure element of their binder-contacting residues as predominantly loop-mediated, predominantly helix-mediated, or as displaying balanced contributions from *β*-strands and loops or from loops and helices, using a threshold of 70% of interface residues (Supplementary Figure S11A).

To characterize the spatial distribution of binder recognition across MHC-fold proteins, contact frequencies were mapped onto the group representatives defined in Section 4.15. When the selected representative did not contain a resolved binder, it was replaced by the structurally closest group member containing one. For each structure containing a resolved binder, MHC *α*_1_-*α*_2_ residues were mapped onto the corresponding group representative using pairwise structural alignment with TM-align [81]. The frequency of binder contacts at each aligned residue position was calculated across the analyzed complexes and stored in the B-factor field of the representative MHC structure, which was then colour-coded for visualization of the recognition footprints (Figure 4B).

## 6 Acknowledgements

We are grateful to the Brazilian Biosciences National Laboratory (LNBio), part of the Brazilian Center for Research in Energy and Materials (CNPEM), a nonprofit organization under the supervision of the Brazilian Ministry of Science, Technology, and Innovation (MCTI), for providing staff and infrastructure for the development of this study. The authors acknowledge the CNPEM Summer Scholarship Program for supporting the participation of the first author in this research. This research used the facilities and the Marvin HPC cluster of the Brazilian Biosciences National Laboratory (LNBio). We thank Joao Victor da Silva Guerra (EDB/CNPEM) for valuable discussions on the algorithm. We also acknowledge the National Laboratory for Scientific Computing (LNCC/MCTI, Brazil) for providing HPC resources of the SDumont supercomputer, which have contributed to the research results reported within this paper. URL: http://sdumont.lncc.br. We also thank Dr. Brian Pierce (University of Maryland) for insightful scientific discussions.

