## Supplementary Material for "Structural Landscape of MHC Complexes through Fold Mining"

### Supplementary Material
### Structural Landscape of MHC Complexes through Fold Mining

A Preprint


**Giovanna Santos de Lima** [1,3], **Paulo Henrique Alves dos Santos** [1], **Samuel Chagas de Assis** [1,2], **Pedro Guilherme B. de S. Nazare** [1], **Carlos Daniel Marques Santos Simões** [1,2], **Elton José Ferreira Chaves** [1], **Gabriel Ernesto Jara** [1], **Rocío Lucía Beatriz Riveros Maidana** [1], and **Helder Veras Ribeiro-Filho** [1,2,*]

[1]Brazilian Biosciences National Laboratory, Brazilian Center for Research in Energy and Materials, Campinas, São Paulo, Brazil

[2]Graduate Program in Pharmaceutical Sciences, Faculty of Pharmaceutical Sciences, University of Campinas, Campinas, São Paulo, Brazil

[3]Department of Fundamental Chemistry, Center for Natural and Exact Sciences, Federal University of Pernambuco, Recife, Pernambuco, Brazil


November 23, 2026

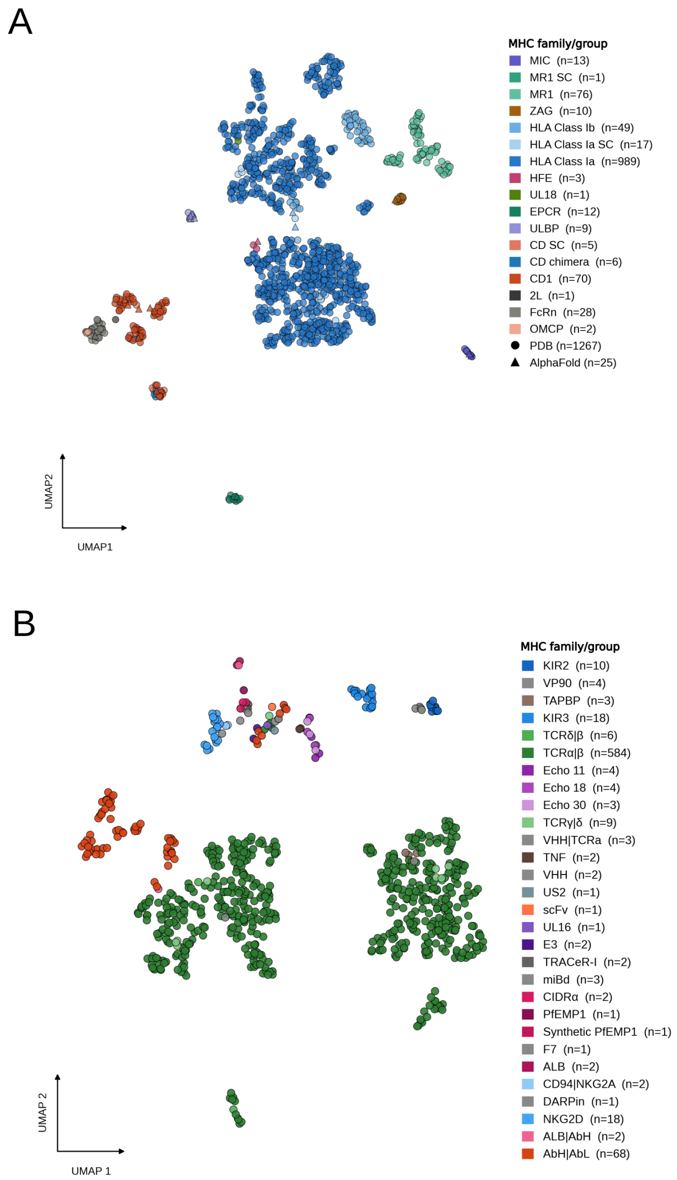


Figure S1: Structural landscape of MHC-fold proteins and their binding partners. (A) Complete two-dimensional UMAP representation of MHC-fold proteins identified in the PDB (n = 1,267) and reviewed UniProt entries retrieved from the AlphaFold Database (AFDB; n = 25), generated from pairwise TM-score similarities of their α1–α2 domains. All structures were included in the UMAP calculation, including single-chain (SC) and chimeric constructs. Structures are colored according to MHC group, with experimentally determined PDB structures represented by circles and AlphaFold models by triangles. Single-chain (SC) and chimeric constructs are distinguished from their corresponding native MHC families. (B) UMAP representation of the structural landscape of MHC-binding partners identified in the PDB, based on pairwise TM-score similarities. Structures are colored according to binder class, with the number of structural instances in each group indicated in the legend.

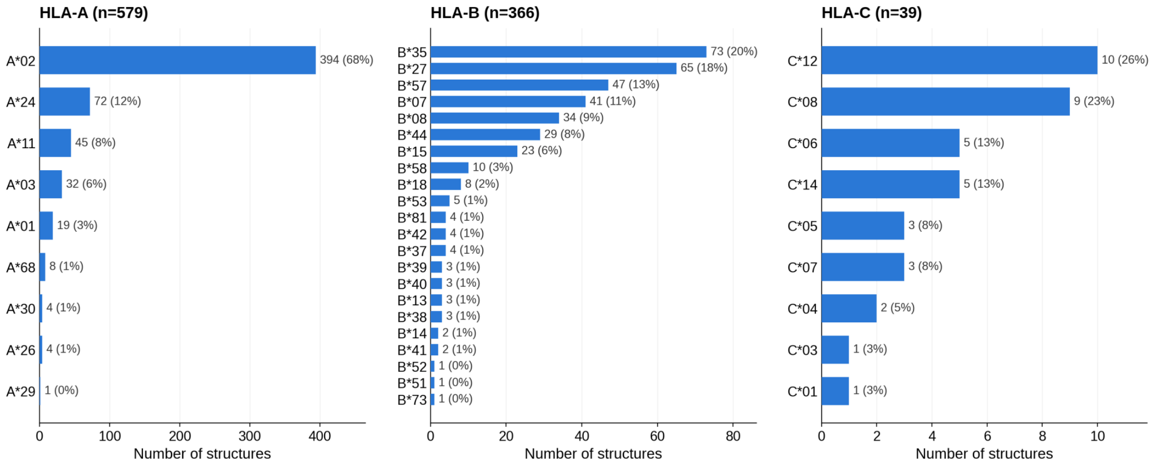


Figure S2: Distribution of classical HLA-I allele groups in the PDB dataset. Number of experimentally determined structures associated with different HLA-A, HLA-B, and HLA-C allele groups. Bars represent the number of structures assigned to each allele group, with absolute counts and corresponding percentages indicated. Percentages were calculated relative to the total number of structures within each HLA locus.

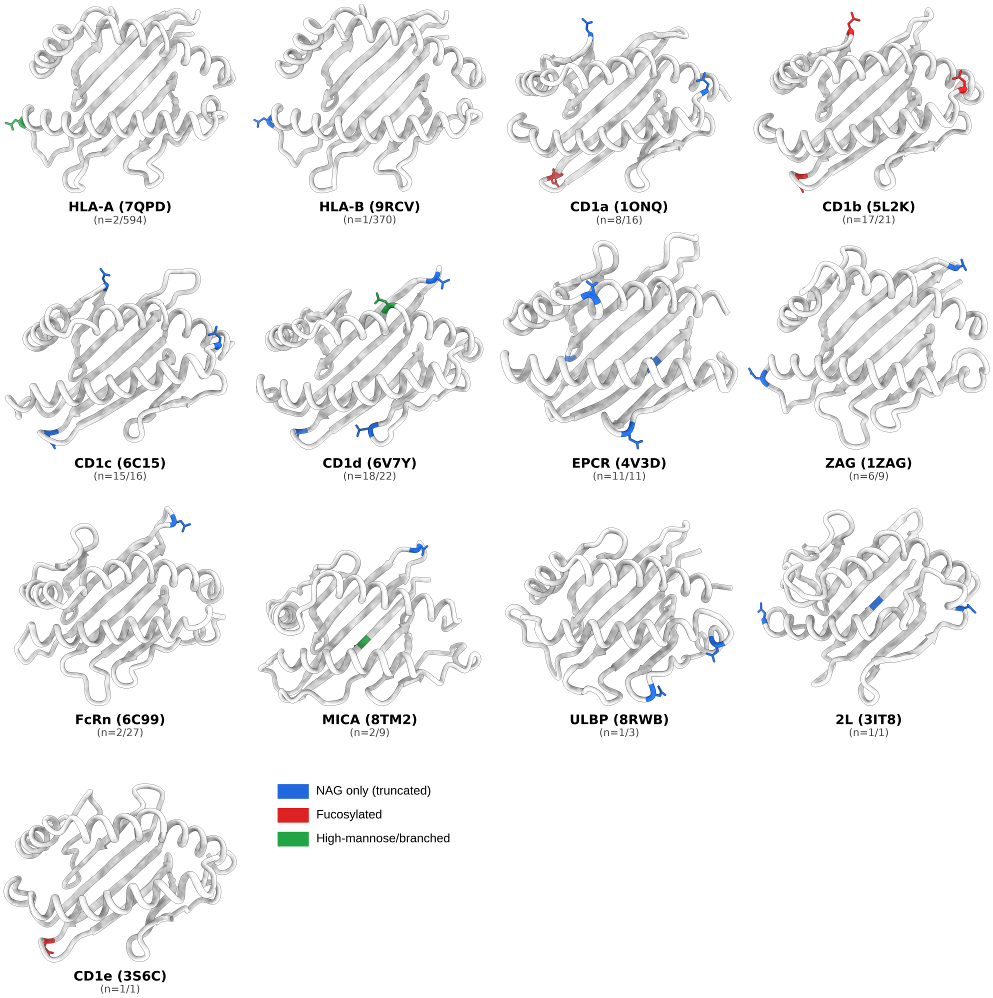


Figure S3: N-linked glycosylation across MHC and MHC-like gene families. One representative structure is shown per gene/family, with confirmed N-glycosylation sites highlighted as sticks and colored by glycan composition: blue, NAG only (truncated); red, fucosylated; green, high-mannose/branched. The PDB ID of the representative structure is given in parentheses below each gene name; n = number of structures with confirmed glycosylation out of the total number of structures for that gene in the dataset.

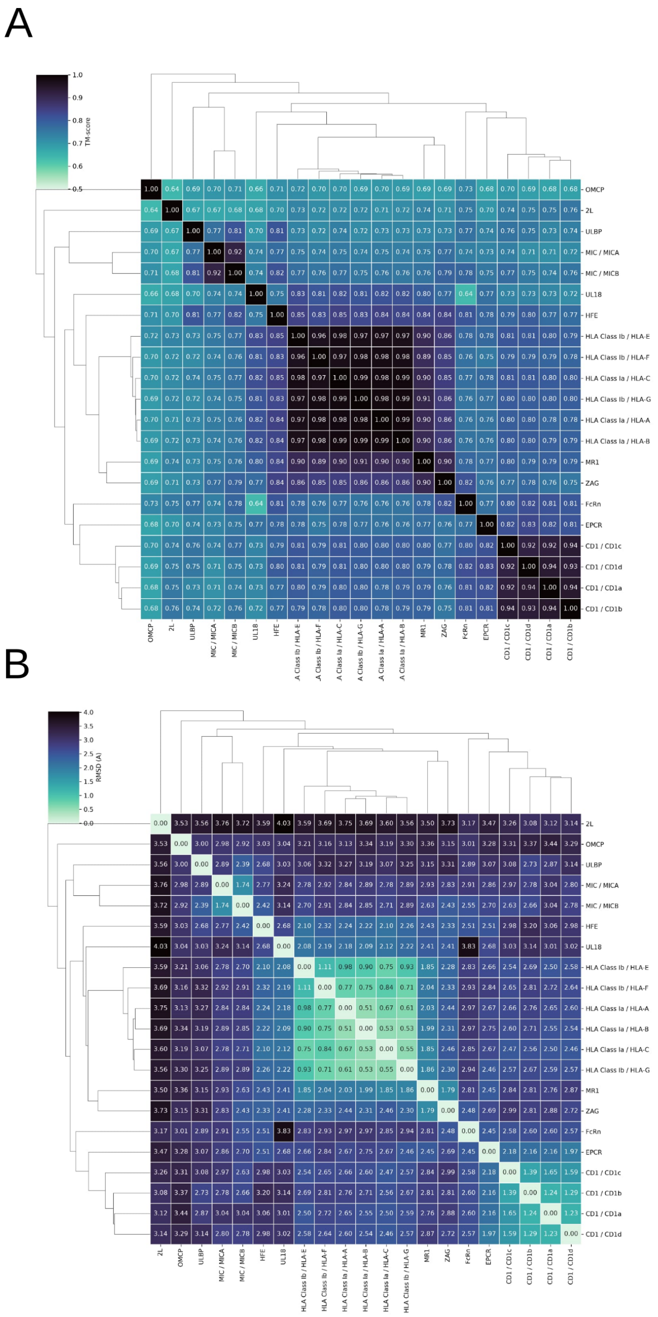


Figure S4: Pairwise structural similarity among representative MHC protein structures. (A) Pairwise TM-score matrix calculated from representative MHC α1–α2 regions across the analyzed protein families using TM-align. Higher TM-scores indicate greater structural similarity, with the corresponding values shown within each cell. Hierarchical clustering based on the pairwise TM-scores is shown by the dendrograms. (B) Pairwise root-mean-square deviation (RMSD) matrix obtained from TM-align superpositions of the same representative MHC α1–α2 regions. RMSD values were calculated over the structurally aligned residue positions identified by TM-align; lower values indicate greater structural similarity. Values are reported in Å and shown within each cell. Hierarchical clustering based on the pairwise RMSD values is shown by the dendrograms.

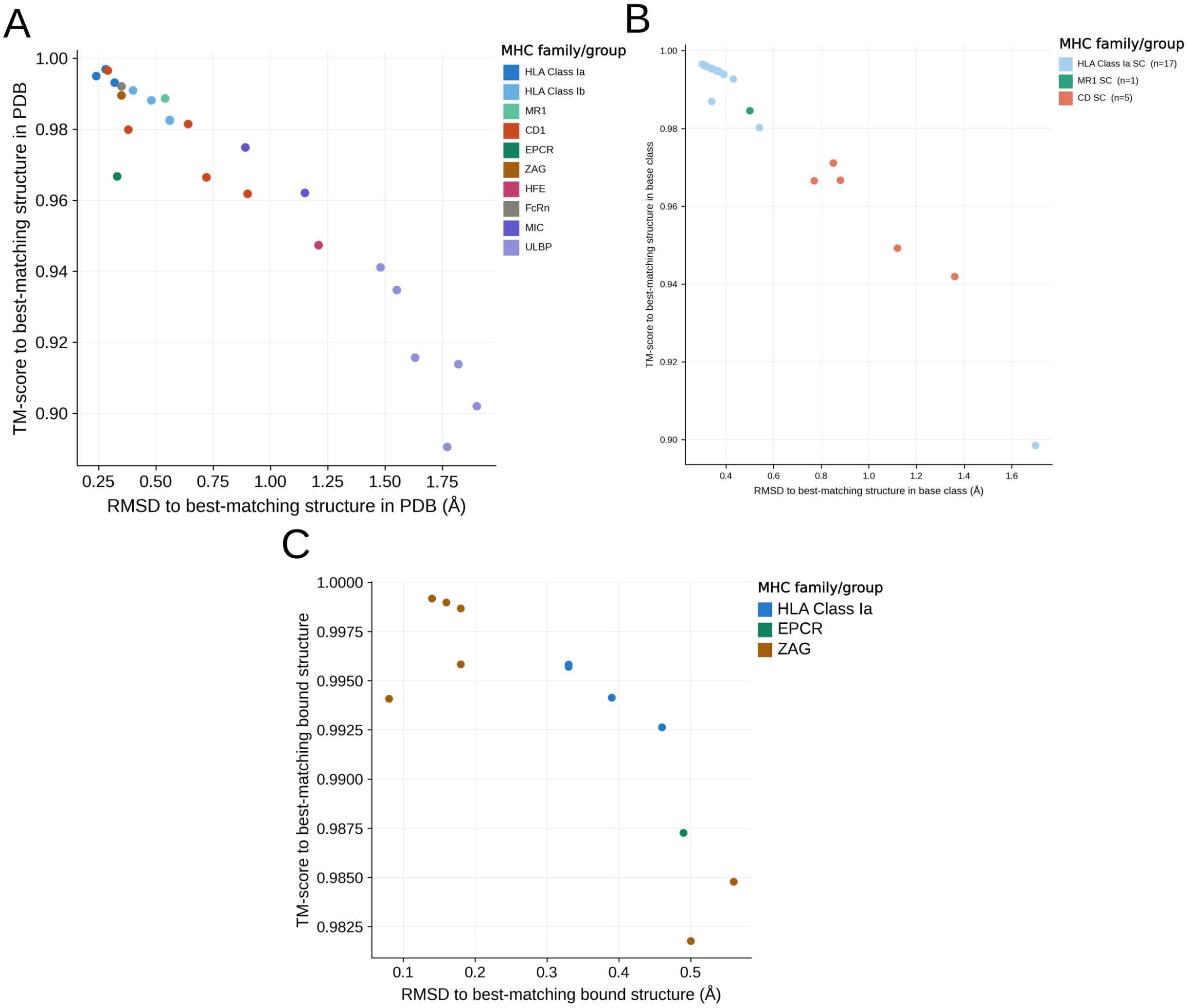


Figure S5: Structural comparison of AlphaFold models and alternative MHC structural states with experimentally determined reference structures. (A) Structural similarity between reviewed AlphaFold Database models of MHC proteins and their best-matching experimentally determined structures in the PDB. Each point represents one AlphaFold model and is colored according to MHC family. The x-axis shows the TM-score to the best-matching PDB structure, and the y-axis shows the corresponding RMSD calculated over the structurally aligned positions. (B) Structural comparison of experimentally determined single-chain MHC constructs with the best-matching structure from their corresponding native MHC family. Single-chain constructs included HLA class Ia (n = 17), MR1 (n = 1), and CD1 (n = 5). TM-scores and RMSD values were obtained from TM-align comparisons. (C) Structural comparison of experimentally determined ligand-free structures with the best-matching ligand-bound structure from the corresponding MHC family. Points are colored according to MHC family, and TM-score and RMSD values were calculated using TM-align over the aligned α1–α2 regions.

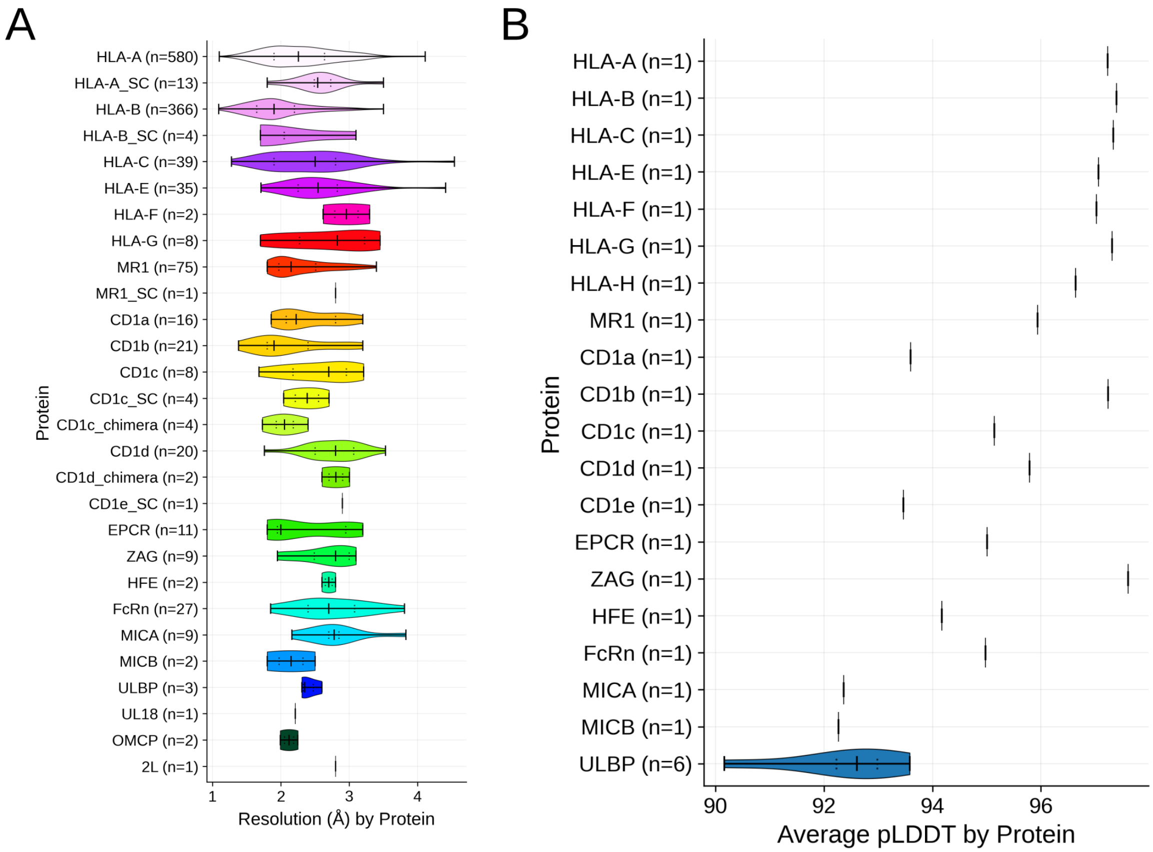


Figure S6: Structural resolution and AlphaFold confidence across MHC proteins. (A) Distribution of experimental resolution across MHC protein structures in the PDB, resolved by protein or family. Single-chain fusion constructs (SC) and chimeric constructs are shown separately where present. Violin plots summarize the distribution of resolution values, with the number of structures analyzed for each group indicated in parentheses. (B) Distribution of mean pLDDT values for reviewed AlphaFold Database models of MHC proteins. The number of models available for each gene or family is indicated in parentheses.

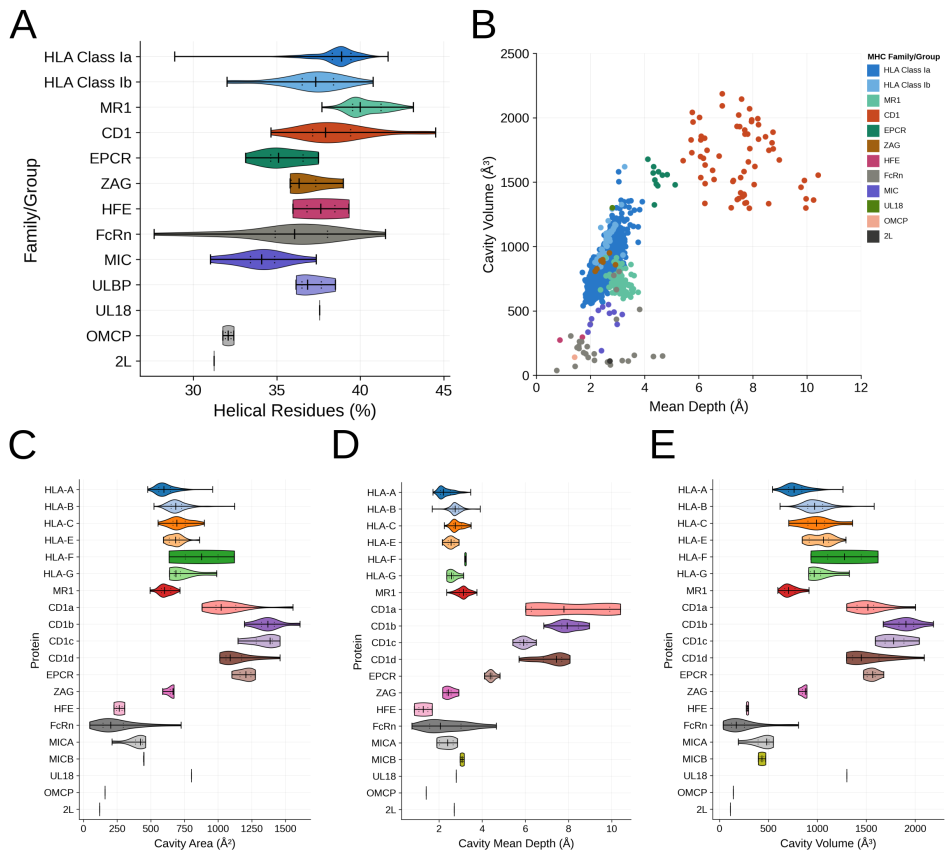


Figure S7: Structural and cavity analysis of MHC proteins. (A) Distribution of the fraction of residues assigned to α-helical secondary structure for each MHC protein family. Helical content was determined using DSSP within the region corresponding to the α1 and α2 domains of classical MHC molecules (see Methods for details). (B) Relationship between cavity mean depth and cavity volume for individual structures, colored according to protein family. Cavity volume and depth were estimated using pyKVFinder (see Methods). (C–E) Distribution of cavity area (C), mean depth (D), and volume (E) resolved at the protein level. Cavity measurements were calculated for the ligand-binding region after removal of the bound ligand, enabling comparison of the intrinsic geometry of binding cavities across classical and non-classical MHC and MHC-like proteins. All analyses were performed after removal of redundant structures.

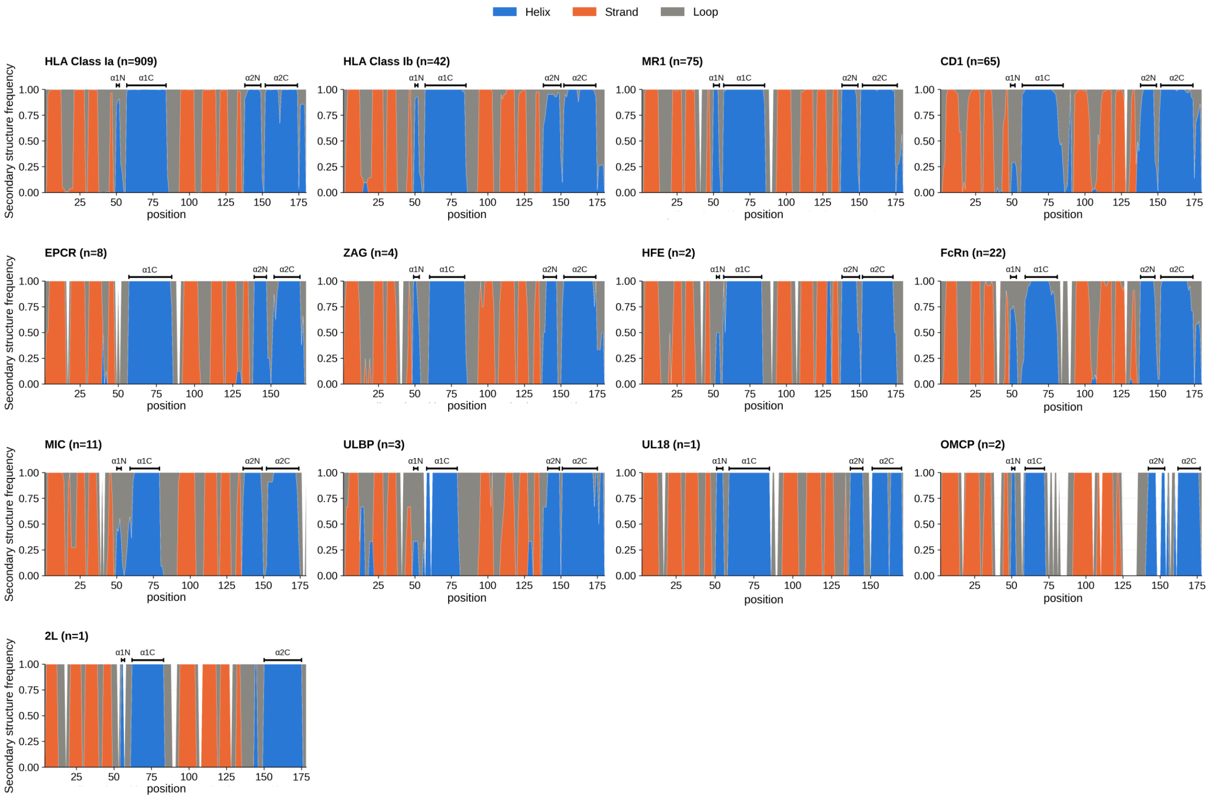


Figure S8: Positional secondary-structure frequencies across MHC-fold protein groups, calculated after structural alignment to the reference HLA-A structure (PDB ID: 3H7B). Stacked profiles indicate the relative frequency of residues assigned to helices (blue), β-strands (orange), or loops (gray) at each aligned position, following the N-to-C direction. The horizontal annotations identify the N-terminal and C-terminal helical subregions of the two canonical MHC groove helices (α1N, α1C, α2N, and α2C), where present.The number of structures analyzed in each group is indicated by n. Analyses were performed after redundancy reduction.

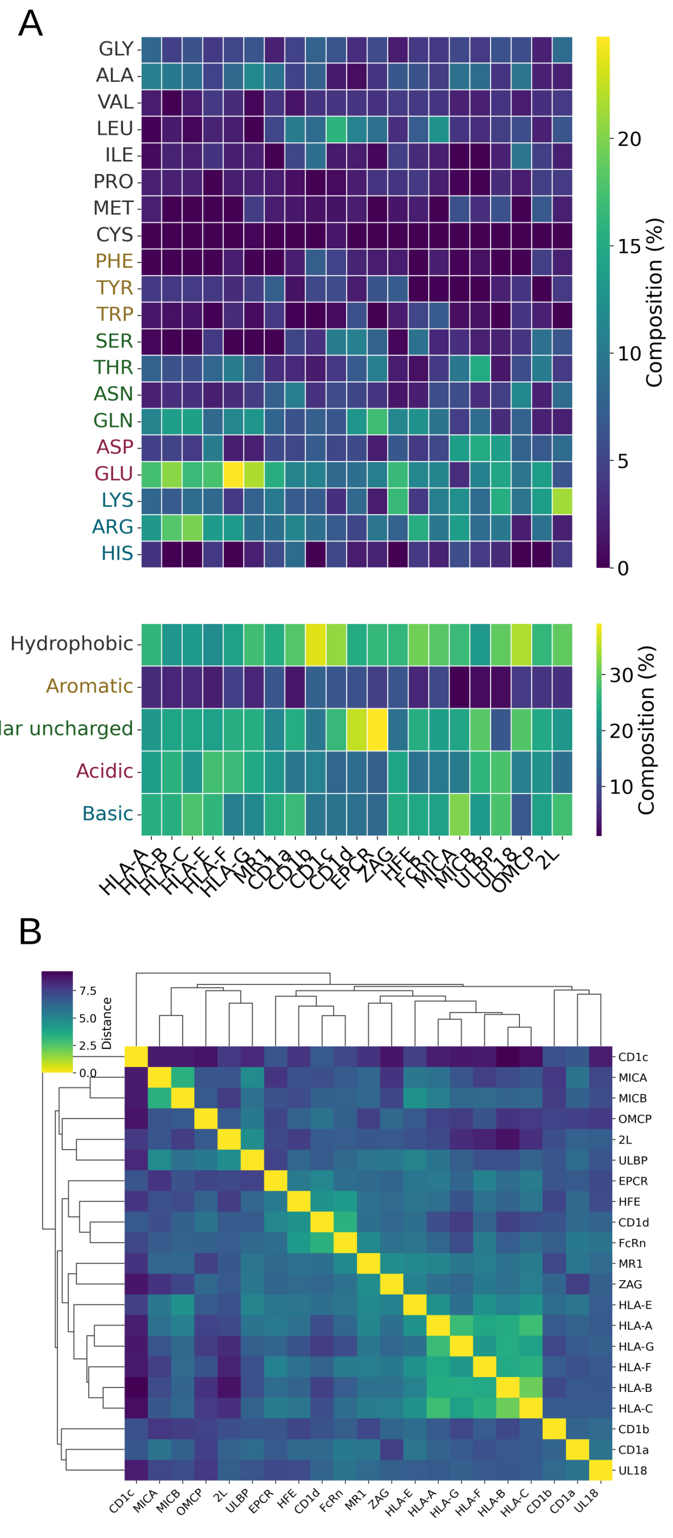


Figure S9: Amino-acid composition and compositional similarity of exposed α1–α2 surfaces across representative MHC proteins. (A) Amino-acid composition of solvent-exposed residues within the α1 and α2 helices for representative MHC proteins, resolved at the protein level. The upper heatmap shows the relative frequency of individual amino acids, whereas the lower heatmap summarizes residues according to the physicochemical classes defined in this study: hydrophobic, aromatic, polar uncharged, acidic, and basic. Frequencies were normalized by the total number of analyzed solvent-exposed helical residues for each protein. (B) Pairwise distance matrix and hierarchical clustering based on the amino-acid composition profiles shown in panel A. Lower distances indicate more similar surface amino-acid composition. All analyses were performed after removal of redundant structures.

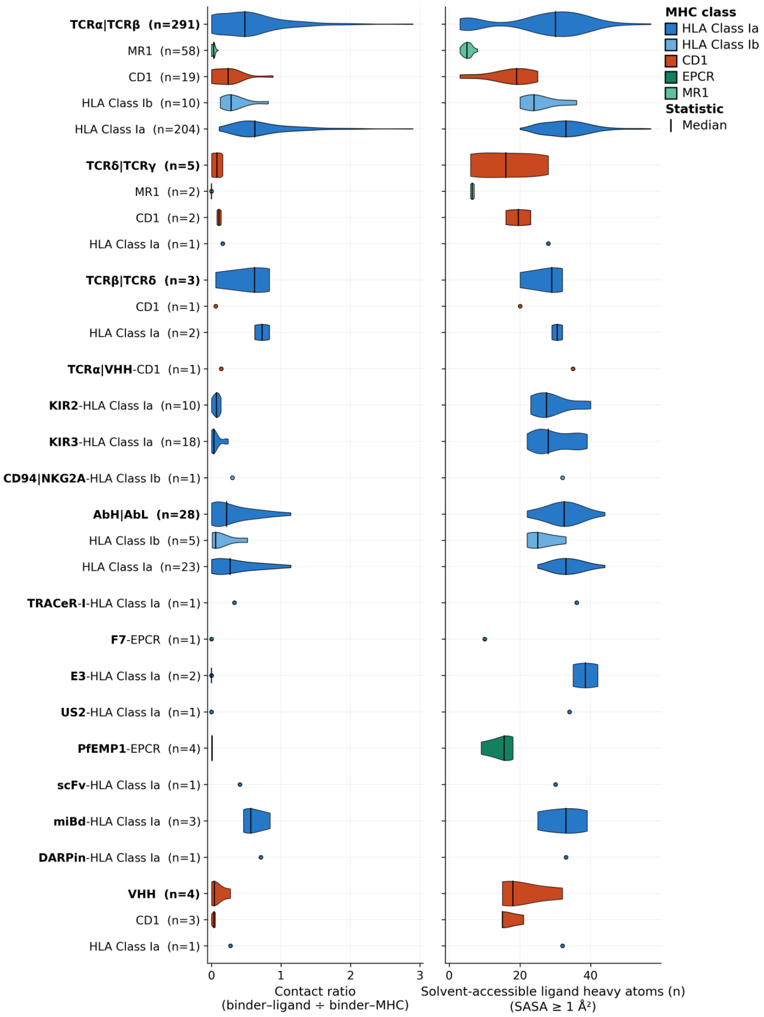


Figure S10: Direct ligand engagement and solvent accessibility across MHC–binder complexes. (A) Distribution of atomic contact ratios between each binder and the MHC-bound ligand relative to contacts between the same binder and the MHC protein (binder–ligand contacts / binder–MHC contacts). Higher ratios indicate a greater relative contribution of direct ligand contacts to the binding interface. (B) Absolute number of solvent-exposed atoms in MHC-bound ligands, considering atoms with solvent-accessible surface area (SASA) >= 1 Å². Complexes are grouped according to binder class and subdivided by MHC family where applicable. Colors indicate the corresponding MHC families, and the number of analyzed complexes (n) is provided in parentheses. Violin plots illustrate the distributions, with solid lines indicating medians, dotted lines indicating quartiles, and white diamonds indicating means. Individual observations are shown for groups with insufficient data to estimate a distribution. All quantitative analyses were performed after removal of redundant structures.

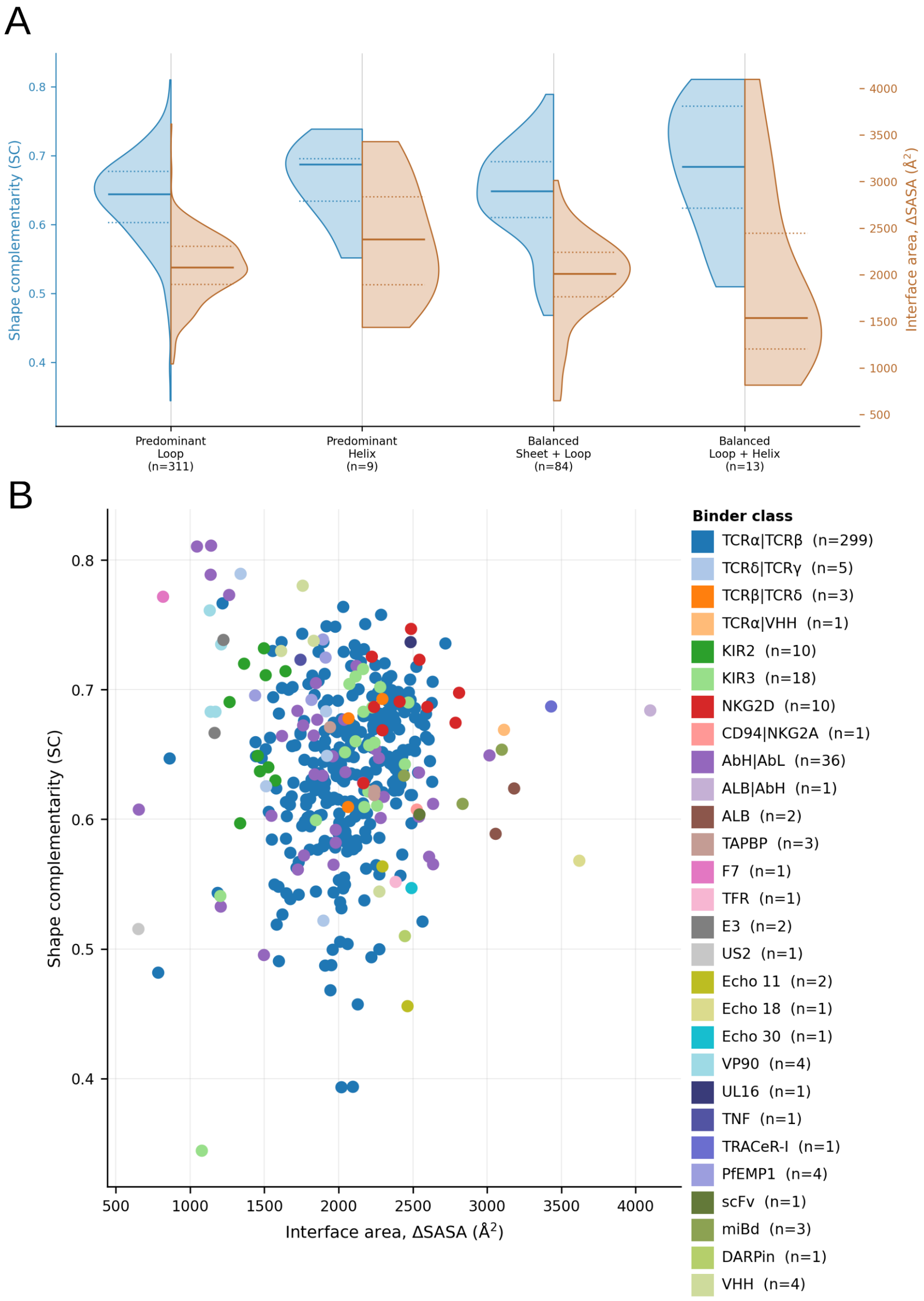


Figure S11: Relationship between binder interface architecture, buried surface area, and shape complementarity across MHC–binder complexes. (A) Distributions of interface shape complementarity (SC, blue, left axis) and buried solvent-accessible surface area (ΔSASA, orange, right axis) across MHC–binder interfaces grouped according to the secondary-structure composition of their contacting residues. Interfaces were classified as predominantly loop-mediated, predominantly helix-mediated, or displaying balanced contributions from β-strands and loops or from loops and helices. Split violin plots show the distributions of both descriptors, with solid lines indicating medians and dotted lines indicating quartiles. The number of analyzed interfaces in each group is indicated in parentheses. (B) Relationship between interface area (ΔSASA) and shape complementarity (SC) across experimentally determined MHC–binder complexes. Each point represents an individual interface, colored according to binder class, with the number of interfaces per class indicated in the legend. No substantial linear correlation was observed between interface area and shape complementarity (Pearson r = 0.03).

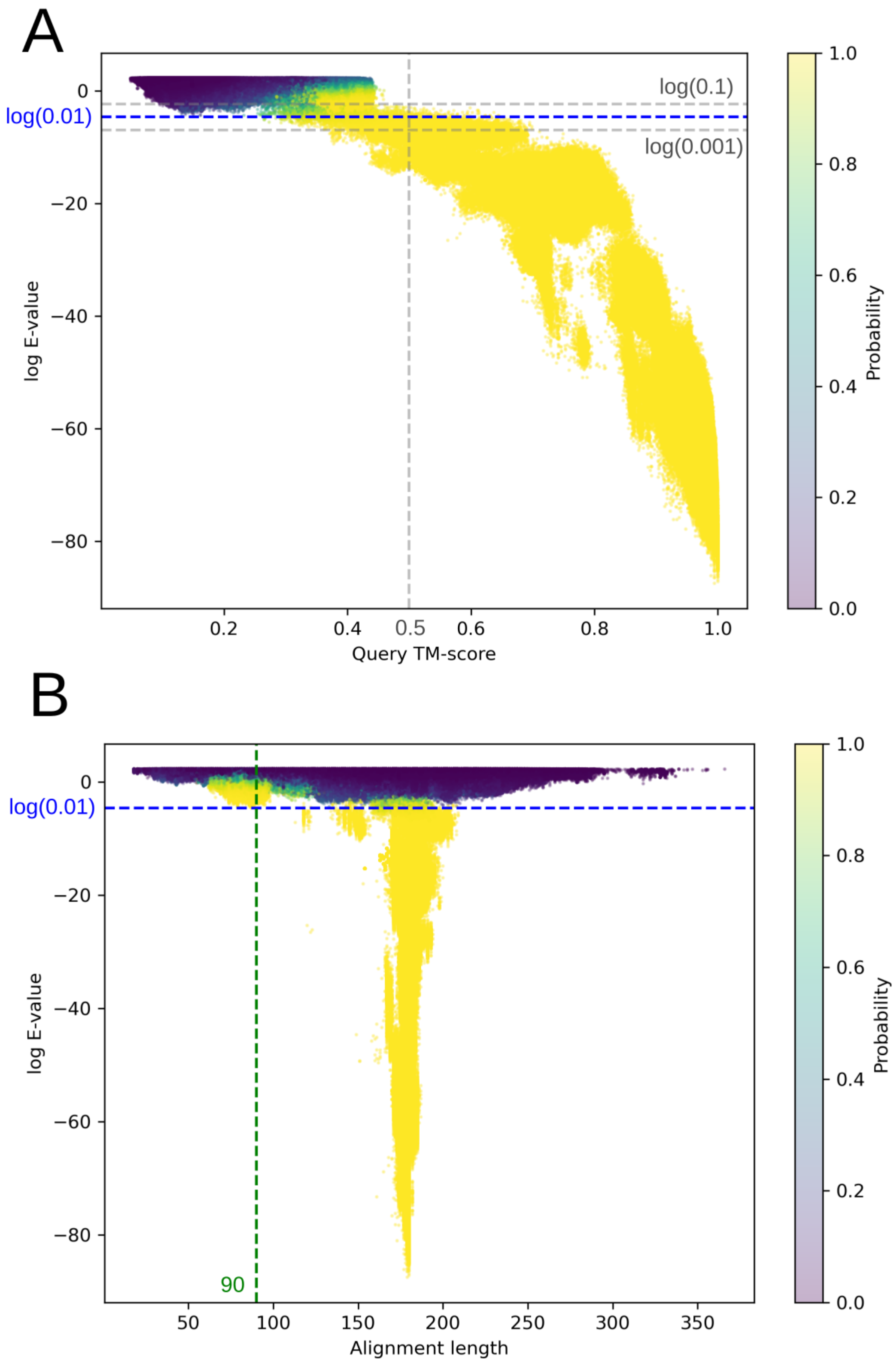


Figure S12: Evaluation of Foldseek criteria for identifying MHC α1–α2 proteins in the PDB. (A) Relationship between the natural-log-transformed Foldseek E-value and query TM-score (qTM-score; TM-score normalized by the query length) for Foldseek hits obtained using the MHC α1–α2 query. Points are colored according to the Foldseek-estimated probability of homology between query and target. The selected E-value cutoff of 0.01 is indicated by the blue dashed line, while E-values of 0.1 and 0.001 are shown as reference thresholds. A qTM-score of 0.5 is indicated by the vertical dashed line. The selected E-value cutoff retains the region containing most high-qTM-score matches while reducing the contribution of weaker structural matches. (B) Relationship between the natural-log-transformed Foldseek E-value and alignment length. The selected E-value cutoff of 0.01 is indicated by the blue dashed line. The alignment-length threshold of 90 residues applied during structural selection is indicated by the green dashed line.

#### References

Table 1: Structures manually excluded prior to download. Structures are ordered alphabetically by PDB ID.

| PDB ID | Reason |
|---|---|
| 1B3J | Very large loop in chain A |
| 1HLA | Structure contains only alpha carbons |
| 1LP9 | Binder from a species outside the project's scope of interest |
| 1LQV | Post-translational modification on a binder chain |
| 2J8U | Binder from a species outside the project's scope of interest |
| 2JCC | Binder from a species outside the project's scope of interest |
| 2UWE | Binder from a species outside the project's scope of interest |
| 3ES6 | Assembly 1 incomplete |
| 3JTC | Post-translational modification on a binder chain |
| 3M17 | Assembly 1 incomplete |
| 3M1B | Assembly 1 incomplete |
| 4LCY | Assembly 1 incomplete |
| 4MNQ | Binder from a species outside the project's scope of interest |
| 4NQC | Assembly 1 incomplete |
| 4NQD | Assembly 1 incomplete |
| 4PJ8 | Gap too narrow to allow modeling |
| 5BJT | Assembly 1 incomplete |
| 5D5M | Assembly 1 incomplete |
| 5D7L | Assembly 1 incomplete |
| 6ENY | Structure lacks side chains, causing the cavity to appear extravasated |
| 6Q3S | Assembly 1 incomplete |
| 6UK2 | Assembly 1 incomplete |
| 6UK4 | Assembly 1 incomplete |
| 7B5F | Gap affecting half of one of the helices; manually compared against other family members (6LA7, 7XXA), only this one has this gap; no associated publication in the PDB |
| 7RNO | Post-translational modification on a binder chain |
| 8DV4 | Assembly 1 incomplete |
| 8EC5 | Gap too large |
| 8I5C | Binder from a species outside the project's scope of interest |
| 8I5D | Binder from a species outside the project's scope of interest |
| 8JHV | No-op entry: does not match any primary assembly from step 1, so it excludes nothing in practice |
| 8TQ6 | Binder from a species outside the project's scope of interest |
| 8WTE | Binder from a species outside the project's scope of interest |
| 8WUL | Binder from a species outside the project's scope of interest |
| 9C96 | Excluded later (not part of the step 2 download blacklist): binder has too many gaps |
| 9CGS | Post-translational modification on the MHC chain |
| 9IKY | Binder from a species outside the project's scope of interest |
| 9LLU | Excluded later (not part of the step 2 download blacklist): TCR chain does not match the one in the associated publication |
| 9O4X | TCR in the wrong assembly |
| 9PBG | TCR in the wrong assembly |
| 9PIX | Binder from a species outside the project's scope of interest |
| 9PKC | Binder from a species outside the project's scope of interest |
| 9PKF | Binder from a species outside the project's scope of interest |
| 9PKV | Binder from a species outside the project's scope of interest |
| 9TF0 | Gap too large |
| 9VSO | Binder from a species outside the project's scope of interest |
| 9Y0T | TCR in the wrong assembly |

Table 2: Non-peptide ligands identified and excluded from the structural dataset. Structures are ordered alphabetically by residue name.

| Residue | Chemical name | Reason |
|---|---|---|
| 1PE | Pentaethylene glycol | PEG analog; crystallization precipitating agent |
| 1PG | 2-(2-{2-[2-(2-Methoxy-ethoxy)-ethoxy]-ethoxy}-ethoxy)-ethanol | PEG analog; crystallization precipitating agent |
| 3NI | Nickel | Metal ion from purification (His-tag) or phasing; experimental artifact |
| ACE | Acetyl | N-terminal acetyl group; chemical modification artifact of the sample |
| ACT | Acetate | Organic solvent; crystallization artifact |
| ACY | Acetic acid | Buffer/crystallization ion; no biological function in the groove |
| B3P | 2-[3-(2-Hydroxy-1,1-dihydroxymethyl-ethylamino)-propylamino]-2-hydroxymethyl-propane-1,3-diol | Crystallization buffer; experimental artifact |
| BGC | D-saccharide (BGC) | Saccharide; glycosylation or crystallization artifact |
| BMA | D-saccharide (BMA) | Saccharide; glycosylation or crystallization artifact |
| BME | Beta-mercaptoethanol | Reducing agent used in sample preparation; experimental artifact |
| BR | Bromide | Phasing derivatizing agent (SAD/MAD); crystallization artifact |
| BTB | 2-[Bis-(2-hydroxy-ethyl)-amino]-2-hydroxymethyl-propane-1,3-diol | Crystallization buffer; experimental artifact |
| CA | Calcium | Buffer/crystallization metal ion; no biological function in the groove |
| CD | Cadmium | Metal ion used in phasing; crystallization artifact |
| CL | Chloride | Buffer/crystallization solution ion; no functional relevance |
| CO | Cobalt | Crystallization or phasing metal ion; experimental artifact |
| CS | Cesium | Phasing metal ion (SAD); crystallization artifact |
| CU | Copper | Crystallization or phasing metal ion; experimental artifact |
| D10 | Decane | Detergent; sample preparation artifact |
| D12 | Dodecane | Detergent; sample preparation artifact |
| EDO | 1,2-Ethanediol | Cryoprotectant; crystallization artifact |
| EDT | {[-(Bis-carboxymethyl-amino)-ethyl]-carboxymethyl-amino}-acetic acid | Chelating agent; purification/sample preparation artifact |
| EMC | Ethyl mercury | Derivatizing reagent for phasing (SAD/MAD); crystallization artifact |
| EPE | 4-(2-Hydroxyethyl)-1-piperazine ethanesulfonic acid | Crystallization buffer; experimental artifact |
| FLC | Citrate | Crystallization precipitating agent; experimental artifact |
| FMT | Formic acid | Buffer/precipitating ion; crystallization artifact |
| FRU | D-saccharide (fructose) | Saccharide; glycosylation or crystallization artifact |
| FUC | L-saccharide (fucose) | Saccharide; glycosylation or crystallization artifact |
| GGG | Glycylglycylglycine (triglycine) | Tripeptide crystallization additive; non-biological, not part of the modeled chains |
| GLC | D-saccharide (glucose) | Saccharide; glycosylation or crystallization artifact |
| GOL | Glycerol | Cryoprotectant; crystallization artifact |
| HEX | Hexane | Organic solvent; crystallization artifact |
| HOH | Water | Solvent molecule; absent in vivo in the MHC groove |
| IMD | Imidazole | Crystallization buffer; experimental artifact |
| IOD | Iodide | Phasing derivatizing agent; crystallization artifact |
| IPA | Isopropyl alcohol | Organic solvent/cryoprotectant; crystallization artifact |
| K | Potassium | Buffer/crystallization solution ion; no functional relevance |
| KZF | 2-(Cyclohexylazaniumyl)ethanesulfonate | CHES buffer (protonated form); crystallization buffer; experimental artifact |
| LMR | (2S)-2-Hydroxybutanedioic acid | Crystallization precipitating agent; presence in the groove attributed to non-specific cavity occupation |

*Continued on next page*

Table 2 (continued)

| Residue | Chemical name | Reason |
|---|---|---|
| MAN | D-saccharide (mannose) | Saccharide; glycosylation or crystallization artifact |
| MES | 2-(N-Morpholino)-ethanesulfonic acid | Crystallization buffer; experimental artifact |
| MG | Magnesium | Buffer/crystallization metal ion; no biological function in the groove |
| MLA | Malonic acid | Crystallization precipitating agent; experimental artifact |
| MLI | Malonate (malonic acid, dianion) | Crystallization precipitating agent; experimental artifact |
| MPO | 3-[N-Morpholino]propane sulfonic acid | Crystallization buffer; experimental artifact |
| NA | Sodium | Buffer/crystallization solution ion; no functional relevance |
| NAG | D-saccharide (NAG) | Saccharide; glycosylation or crystallization artifact |
| NDG | D-saccharide (NDG) | N-glycosylation saccharide; post-translational modification artifact |
| NHE | 2-[N-Cyclohexylamino]ethane sulfonic acid | CHES buffer; experimental artifact |
| NI | Nickel | Metal ion from purification (His-tag) or phasing; experimental artifact |
| NO3 | Nitrate | Buffer/crystallization ion; no biological function in the groove |
| P4G | 1-Ethoxy-2-(2-ethoxyethoxy)ethane | PEG analog; crystallization precipitating agent |
| PEG | Di(hydroxyethyl)ether | Crystallization precipitating agent; no biological function |
| PG4 | Tetraethylene glycol | PEG analog; crystallization precipitating agent |
| PGE | Triethylene glycol | Cryoprotectant; crystallization artifact |
| PO4 | Phosphate | Crystallization buffer ion; no biological function in the groove |
| R16 | Hexadecane | Long-chain aliphatic hydrocarbon; detergent artifact |
| SCN | Thiocyanate | Crystallization solution salt; experimental artifact |
| SO4 | Sulfate | Contaminant/buffer; no biological function in the groove |
| TAM | Tris(hydroxyethyl)aminomethane | Crystallization buffer; experimental artifact |
| TRS | 2-Amino-2-hydroxymethyl-propane-1,3-diol | Crystallization buffer; experimental artifact |
| TWT | Docosane | Long-chain aliphatic hydrocarbon; detergent artifact |
| UND | Undecane | Long-chain aliphatic hydrocarbon; detergent artifact |
| Y01 | Cholesterol hemisuccinate | Detergent/lipid additive used in membrane-protein crystallization; experimental artifact |
| ZN | Zinc | Crystallization or phasing metal ion; experimental artifact |

Table 3: Binder chains manually removed from the structures. The structures were retained in the dataset. Entries are ordered alphabetically by PDB ID.

| PDB ID | Chain | Reason |
|---|---|---|
| 28IL | C | $\beta$2-microglobulin chain assigned as binder |
| 7DZM | B | $\beta$2-microglobulin chain assigned as binder |
| 7NDT | F | Copy from the asymmetric unit (chain E) |
| 7NUI | B | $\beta$2-microglobulin chain assigned as binder |
| 8DV3 | B | $\beta$2-microglobulin chain assigned as binder |
| 9F13 | D | $\beta$2-microglobulin chain assigned as binder |
| 9NW9 | B | $\beta$2-microglobulin chain assigned as binder |
| 9O06 | F | Copy of TCR from another MHC copy |
| 9Y0T | D | $\beta$2-microglobulin chain assigned as binder |
| 9Y0U | F | Copy of TCR from another MHC copy |
| 9Y0W | F | Copy of TCR from another MHC copy |
